# Cryo-EM structure of the folded-back state of human β-cardiac myosin*

**DOI:** 10.1101/2023.04.15.536999

**Authors:** Alessandro Grinzato, Daniel Auguin, Carlos Kikuti, Neha Nandwani, Dihia Moussaoui, Divya Pathak, Eaazhisai Kandiah, Kathleen M. Ruppel, James A. Spudich, Anne Houdusse, Julien Robert-Paganin

## Abstract

During normal levels of exertion, many cardiac muscle myosin heads are sequestered in an off-state even during systolic contraction to save energy and for precise regulation. They can be converted to an on-state when exertion is increased. Hypercontractility caused by hypertrophic cardiomyopathy (HCM) myosin mutations is often the result of shifting the equilibrium toward more heads in the on-state. The off-state is equated with a folded-back structure known as the interacting head motif (IHM), which is a regulatory feature of all muscle myosins and class-2 non-muscle myosins. We report here the human β-cardiac myosin IHM structure to 3.6 Å resolution. The structure shows that the interfaces are hot spots of HCM mutations and reveals details of the significant interactions. Importantly, the structures of cardiac and smooth muscle myosin IHMs are dramatically different. This challenges the concept that the IHM structure is conserved in all muscle types and opens new perspectives in the understanding of muscle physiology. The cardiac IHM structure has been the missing puzzle piece to fully understand the development of inherited cardiomyopathies. This work will pave the way for the development of new molecules able to stabilize or destabilize the IHM in a personalized medicine approach.

*\*This manuscript was submitted to Nature Communications in August 2022 and dealt efficiently by the editors. All reviewers received this version of the manuscript before 9^208^ August 2022. They also received coordinates and maps of our high resolution structure on the 18^208^ August 2022. Due to slowness of at least one reviewer, this contribution was delayed for acceptance by Nature Communications and we are now depositing in bioRxiv the originally submitted version written in July 2022 for everyone to see. Indeed, two bioRxiv contributions at lower resolution but adding similar concepts on thick filament regulation were deposited this week in bioRxiv, one of the contributions having had access to our coordinates.*

*We hope that our data at high resolution will be helpful for all readers that appreciate that high resolution information is required to build accurate atomic models and discuss implications for sarcomere regulation and the effects of cardiomyopathy mutations on heart muscle function.*

---

Muscle contraction depends on myosins from class 2. In the sarcomeres, they associate in filaments: the so-called thick filaments that interact with the thin filaments of filamentous actin to produce force during contraction (reviewed by ^1^). Muscle contraction is energy consuming and fueled by ATP, and skeletal muscles represent between 30 and 40% of the body mass for an adult human^2^. In order to avoid energy loss between contractions, the myosin heads adopt a super-relaxed state (SRX) characterized by a low ATP turnover^3–5^, promoting energy saving^3, 5^. This functional state is present in all muscle fibers including skeletal^3^; cardiac^5, 6^ and smooth^7^ muscles.

While the structural basis underlying force production by myosins has been intensively studied (reviewed by ^1^), the structural basis of the SRX remains poorly understood. From early studies on smooth muscle myosins (SmMyo2), the SRX has been linked to the asymmetrical **I**nteracting **H**eads **M**otif (IHM) where the two heads of a myosin molecule fold back onto their own coiled-coil tail^8, 9^. The IHM was further recognized from structural studies of human cardiac myo2^10^, non-muscle myo2^11^ and muscles from animals such as sea sponges and jellyfish^11^, worms, insects, arachnids, mollusks, fish and human^12^. From the low resolution examined (>2 nm), all types of muscle myo2 IHM look alike and this motif has been assumed to be conserved^11^. Three higher resolution cryoEM structures of the smooth muscle myosin (SmMyo2) IHM have been reported at resolutions of 6 Å^13^, 4.3 Å ^14^ and 3.4 Å ^15^. These structures reveal the intra molecular interactions within the dimer (IHM interfaces) and the different hinges of flexibility required for the stabilization of this sequestered state for smooth muscle myosin.

The role of myosin filament-based regulation to control the time course and strength of cardiac contraction has been demonstrated ^16^. Hypertrophic cardiomyopathy (HCM)-causing point mutations in human β-cardiac myosin, the motor that drives heart contraction, have been reported to destabilize the SRX and increase the number of heads available to engage force production, leading to hypercontractility (eg.,^17^). Many of these HCM mutations are located on a flat surface called the “Mesa” and have been proposed to lead to the destabilization of the IHM interfaces, thus increasing force^18–20^. Other mutations may affect the stability of diverse conformational states of the motor leading to different ways of modulating the force produced^21^. Interestingly, the recent SmMyo2 IHM structures challenges the Mesa hypothesis^14^ in that these structures are significantly different from the human β-cardiac homology models^20^. Moreover, the activator *Omecamtiv mecarbil* (OM, in phase 3 clinical trials) and the inhibitor *Mavacamten* (Mava, approved by the FDA for treatment of HCM) have been reported to destabilize^22^ and stabilize^23^ respectively the sequestered state. The sequestered state of β-cardiac myosin has been described at 28 Å resolution^10^, but that is insufficient to describe precisely the IHM interfaces. Together, these results emphasize the urgent need for a high-resolution structure of the cardiac myosin IHM to understand the development of inherited cardiac diseases, and to treat these diseases via the development of new modulators acting on the stability of the sequestered state.

Here, we present the high-resolution structure of the human β-cardiac myosin IHM solved at 3.6 Å resolution, with a highest resolution in the heads of 3.2 Å. Importantly, the comparison with SmMyo2 IHM reveals some major differences in the interfaces and the hinges in the lever arm and the coiled-coil region, challenging the claims that the IHM are conserved amongst the different myosins 2 and that SmMyo2 IHM structure would have been a good model to discuss the cardiomyopathy mutations. Altogether, our results provide an accurate framework for studying the effects of HCM mutations, and for understanding small molecule modulator effects on the stability of the sequestered state. Finally, the structure reveals how the differences between the cardiac and the smooth IHM may be an adaptation to fit different muscle functions and organization.

## Structure of the cardiac myosin IHM

We solved the near-atomic resolution IHM structure of human β-cardiac myosin-2 via single-particle cryo-EM using a native uncrosslinked protein preparation with ATP bound (see Methods and **Extended data Fig. 1**). In SmMyo2 but not Cardiac myo2, a distal coiled-coil region (residues 1410-1625) adds to the stability of the sequestered, auto-inhibited state in addition to the S2 coiled-coil ^10^(**Extended data Fig. 2**). The dimeric cardiac myosin construct we used comprised two heads (human β-cardiac motor domain followed by the lever arm with human β-cardiac essential (ELC) and regulatory (RLC) light chains bound to the target HC sequences IQ1 and IQ2 respectively) and 15 heptads of the proximal S2 coiled-coil. The IHM classes used for the final 3D reconstruction represented only a portion of the picked particles (∼200 000 particles over 0.5 million). We thus obtained the high-resolution structure of the sequestered off-state (IHM) of human β-cardiac myosin at a global resolution of 3.6 Å (map 1) (**Extended data Table 1, Extended data Fig. 1, Fig. 1a**). The IHM motif was immediately recognizable in the cryo-EM map, with the so-called blocked and free heads (BH and FH respectively), the lever arm and the proximal S2 coiled-coil. In map 1, the head regions correspond to the highest resolution and the lever arm/coiled coil to the lowest resolution (**Fig. 1a**, **Extended data Fig. 1, Extended data Movie 1**). By performing a focused refinement on the head/head region, we improved the map for the motor domains to a highest resolution of 3.2 Å (map 2, **Fig. 1b**, **Extended data Movie 2**), with clear indications for the side chains (**Fig. 1c**). The nucleotide bound in each head is clearly ADP and P_i_, which are trapped after hydrolysis of ATP (**Fig. 1c, 1d**). Interestingly, despite the high resolution of the map for both motor domains, some side chains are not visualized although they are likely involved in intra-molecular interactions stabilizing auto-inhibition (**Fig. 1e**). Flexibility exists indeed in the formation of these interactions, as shown also from the movies of the 2D classes (**Extended data Fig. 1f**). The lability of these interactions is an important feature of the IHM stabilization that is required so that the head can be readily activated when they are needed for contraction. We thus computed realistic interfaces with side chain interactions using molecular dynamics (see Methods). Finally, two essential regions of regulation, the N-term extensions of the ELC and RLC were partially rebuilt in density (**Fig. 1f**). In contrast to the SmMyo2 IHM structures, the density of the FH is equivalent to that of the BH in our maps, indicating that both heads are equally stabilized in the cardiac IHM (**Extended data Fig. 1**).

**Figure 1.**
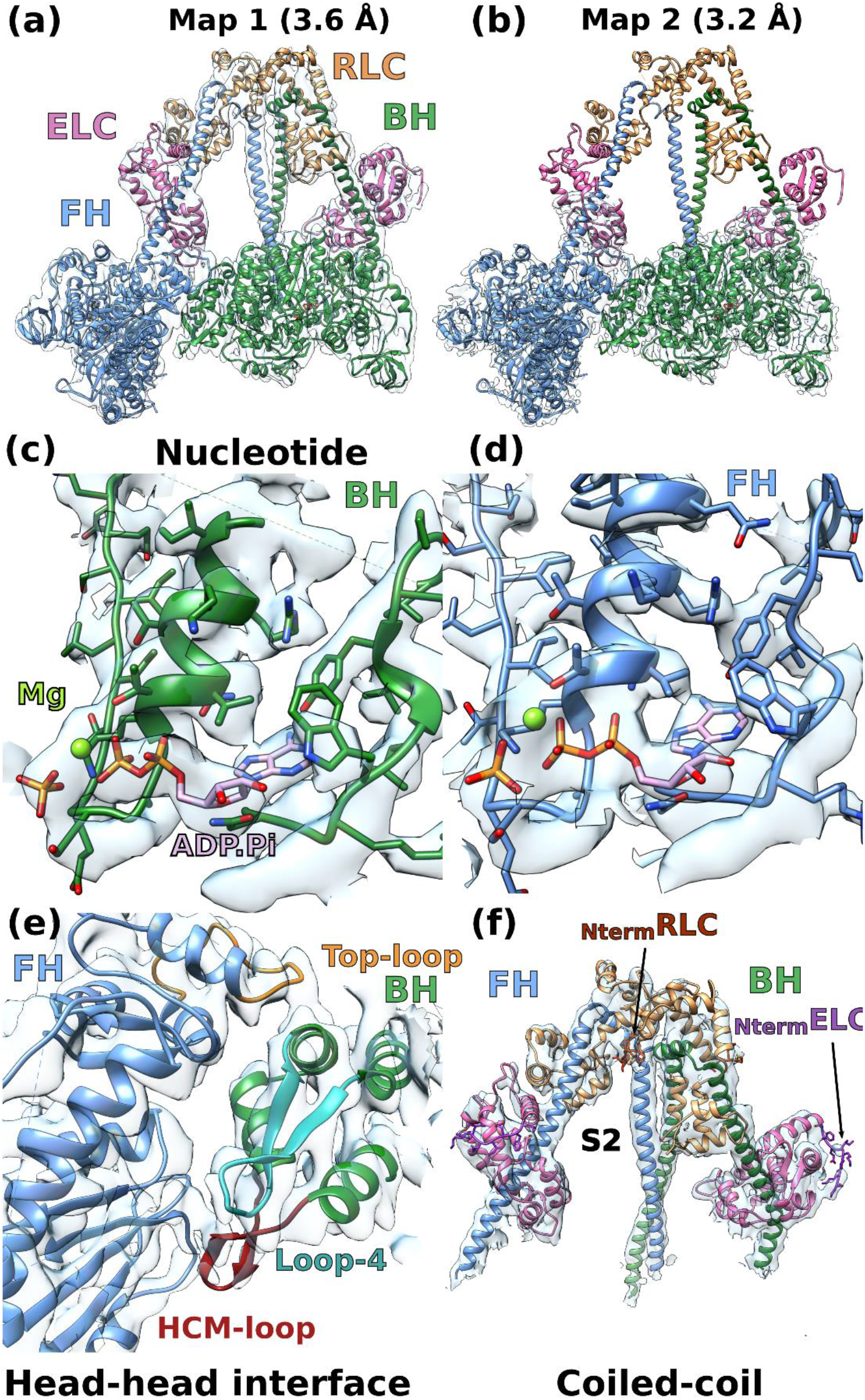
– CryoEM density map of the IHM of β-cardiac myosin. **(a)** Complete map of the interacting head motif (IHM) at a resolution of 3.6 Å (map 1). **(b)** Map masked on the two heads improved the resolution at 3.2 Å (map 2). **(c)** and **(d)** map 2 allows to see the ADP, the hydrolyzed phosphate and the magnesium. The side chains of the residues in the active site are well defined in density. **(e)** Interface between the two heads of the IHM (map 1). **(f)** Coiled-coils, light-chains and N-terminal extensions are seen without ambiguity in map 1, the N-terminal extension (Nterm) are shown by an arrow.

In the cardiac IHM, both heads have the lever arm primed and their motor domains are highly superimposable (rmsd 0.7 Å) (**Extended data Fig. 3a**). Their structural state corresponds to a classical pre-powerstroke (PPS) state (rmsd ∼0.84 Å with 5N6A,^24^), defined by the structure of the motor domain from the N-terminus through the Converter ending at the pliant region (**Extended data Fig. 3b, 3c**). The major and only difference is the larger kink in the pliant region^25^ of the BH lever arm that alters the ELC/Converter interface (**Extended data Fig. 3d, 3e, 3f**) and the so-called musical chairs (**Extended data Fig. 3d**), a network of labile electrostatic interactions involved in the control of the lever arm dynamics at this interface^21^. Our results demonstrate that there is no significant conformational change of the Converter orientation upon formation of the sequestered state from the PPS state. The closed nucleotide pocket and the myosin inner cleft do not change either (**Extended data Fig. 3g**), as suggested by previous spectroscopy experiments ^26^. Despite its overall asymmetry, the IHM has two motor domains essentially in the same conformation.

We compared the high resolution cardiac IHM structure with the three previous homology models (PDB code 5TBY^27^; MS03^28^; MA1^21^) generated using distinct approaches. Since MS03 and 5TBY share similar features, we used only MS03 in the illustration (**Extended data Fig. 4a**). As expected, these models could only approximate the overall position and interface of the heads. The exact positioning of the structural elements engaged in this interface were difficult to predict in the absence of high-resolution structural information (**Extended data Fig. 4b**). The models could not list correctly the residues on the BH head that could be close enough to interact with the S2 coiled-coil (**Extended data Fig. 4c**). The major differences between the models and the structure reside in **the lever arm,** which gives rise to the overall asymmetry in the structure (**Extended data Fig. 4d, 4e**). Indeed, higher resolution was absolutely required to describe correctly the hinges of flexibility.

The MA1 model correctly positioned the BH and FH Converters by keeping their position as found in the single head PPS state (5N6A^24^) ((**Extended data Fig. 4f**), but the lever arms were modelled as straight, which is true only for the FH (**Extended data Fig. 4e**). The other models positioned the Converter in a more primed position for both heads by proposing that the Relay orientation would change to further prime the Converter by 26° (**Extended data Fig. 4f**). In fact, in MS03, large kinks are proposed for the pliant region of both heads (**Extended data Fig. 4d, 4e**). The high resolution IHM structure now reveals that the Converter position is <u>not</u> modified compared to what is found in an active head but the priming of the lever arm occurs only in the BH head and comes from the modification in the pliant region (**Extended data Fig. 4c**). The MS03 model had detected the requirement of kinks in the heads and thus had proposed that the heads would adopt a so-called “pre-prestroke” conformation, with a Converter more primed than the conventional prestroke^20^. This ‘pre-prestroke’ concept implied that slower release of products could result from these conformational changes. The high-resolution IHM structure now reveals that the slow product release (SRX) is not a result of a change in the motor conformation. In the IHM structure, both heads are in the same conventional prestroke conformation and it is the interactions between them that slows their activity.

## The interactions stabilizing the IHM

The structure reveals five main stabilizing regions within the asymmetrical IHM: the head-head interface, the coiled-coil/BH, the RLC/RLC and the BH and FH ELC/RLC interfaces (**Fig. 2**, **Extended data Table 2**). **Extended data Fig. 5** defines the structural elements involved in these interfaces. Both the head-head and the coiled-coil/BH interfaces involve part of a large and flat surface of myosin heads called the Mesa (**Fig. 2a**; **Extended data Fig. 6**), which was originally defined as a large and relatively flat surface containing highly conserved residues among β-cardiac myosins across species, which are hot-spots for cardiomyopathy mutations, and which was predicted to be a docking site for other protein elements that would put the myosin head in an OFF state^18^. More specifically, the mesa was proposed to be essential in the interaction with S2, and possibly MyBPC or other components of the thick filament^20^. Our high-resolution human b-cardiac myosin IHM structure now provides a precise definition of the Mesa residues of the BH head that interact with the S2 coiled-coil, and indicates which Mesa residues of the FH head are involved in a distinct stabilization surface. Mutation of some Mesa residues, such as R453, can thus affect two stabilization surfaces if both the BH and FH heads possess the mutation.

**Figure 2.**
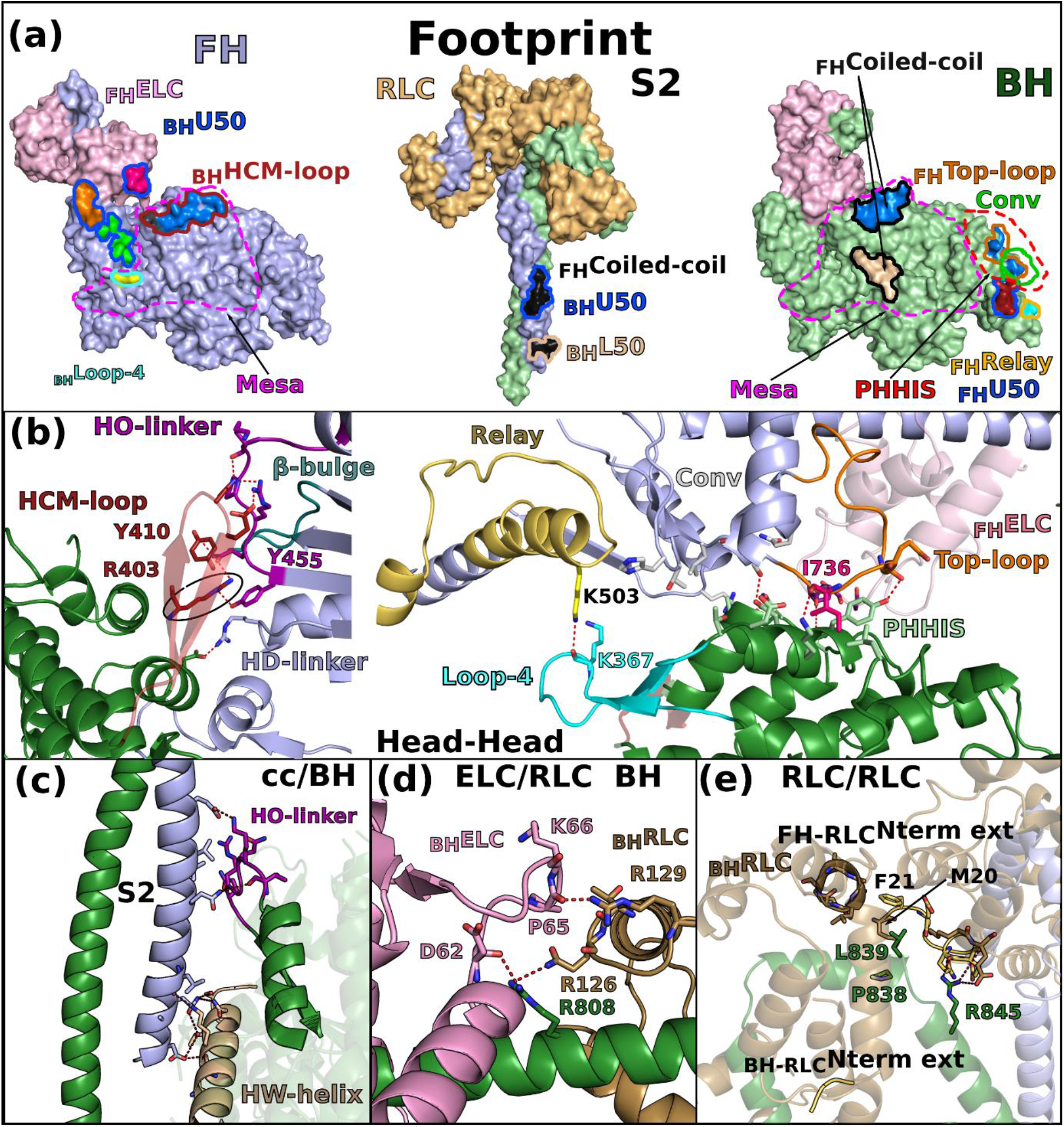
– The IHM is stabilized by multiple interfaces. **(a)** Footprint of the interfaces stabilizing the sequestered state. The subdomains involved in the interaction are colored and the contours of interface surfaces indicate which region of the IHM is involved in recognizing this surface : L50 (wheat); U50 (marine blue); Relay (yellow); Converter (green); coiled-coil (black); HCM-loop (dark red); Loop-4 (cyan). The large flat surface called “Mesa” is delimited by purple dotted lines. **(b)** Cartoon representation of the head-head interface. Key-residues involved in the interaction are represented as stick. The two views on the left and on the right represent the interaction involving the HCM-loop and the interactions mediated by the primary head-head interaction site (PHHIS, in red dotted lines) and Loop-4 respectively. The key-interactions mediated by the HCM-loop through R403 are contoured in a black circle. The connectors and elements involved in the interface are shown and named in **Extended data Fig. 5**. **(c)** Coiled-coil/BH interface. **(d)** ELC/RLC interface on the BH, the differences with the FH are illustrated in **Extended data Fig. 3**. **(e)** RLC/RLC interface involving the N-terminal extension of the RLC from the FH (_FH-RLC_Nterm ext).

The **head-head interface is formed by close interactions between** the Mesa of the FH head with two BH actin-binding elements: loop-4 and HCM-loop as well as residues on the surface of the U50 subdomain previously designated as the primary head-head interaction site (PHHIS) (**Fig. 2a**, **Extended data Fig. 6, Extended data Table 2**). The structural elements of the Mesa in the FH head are the Relay helix (a connector essential for allosteric communication between the active site and the lever arm)^1^, the β-bulge and HO-linker, two connectors of the Transducer^1, 29^ previously reported to influence myosin transient kinetics and force production (**Extended data Table 2**). Interestingly, we find that the HD-linker, an element of the Transducer adjacent to the HO-linker, is also part of the Mesa and interacts with the BH PHHIS (**Fig. 2b**). In addition, the Converter and the ELC loop 3 also interact with the BH PHHIS. This interface includes in particular the specific _Converter_Top-loop, whose flexibility and conformation greatly depends on the sequence of the Converter and its interactions with the ELC^21^ (**Fig. 2b**). Three hallmarks of the head-head interface can be reported (**Extended data Table 2**): **(i)** a high number of electrostatic interactions; **(ii)** the central hydrophobic lock-and-key interaction involving the FH _Top-loop_I736; **(iii)** the importance of the BH _HCM-Loop_R403 that is involved in electrostatic interactions and a cation-π stacking with _HO-linker_Y455, a residue of the FH Mesa (**Fig. 2b**). The BH loop-4 establishes few interactions with the FH _Relay_K503 (**Fig. 2b**). The **coiled-coil (cc)/BH head** interface involves the HO-linker and the _L50_HW helix, comprising some residues of the Mesa of the BH, which both interact with electrostatic surfaces of the S2 coiled-coil while some hydrophobic residues consolidate the interactions (**Fig. 2c**).

**The hinges and interfaces within the lever arm** participate in the stabilization as well as the asymmetric orientation of the two heads of the IHM. The role of flexible hinges in the lever arm for the formation of the IHM had been anticipated^30^ but they had never been described precisely. Here the model built in clear electron density demonstrates that asymmetry in the various hinges within the lever arm are positioned **(i)** at the **pliant region**, located at the end of the Converter (residues 778-782) where the main helix of the lever arm is kinked by ∼55° in BH but remains straight in FH (**Extended data Fig. 3a**); **(ii)** at the **ELC/RLC** interface since differences between the two heads in the ELC loop1 conformation and in the nearby HC kink near R808 promote different electrostatic and apolar interactions at the ELC/RLC interface (**Fig. 2d**, **Extended data Fig. 7a, 7b**); **(iii) within the RLC lobes** in which we find differences between the two heads in the positioning of the HC helices in particular for the bulky residues W816 near the inter-lobe RLC linker as well as W829 and Y823 within the N-terminal lobe of the RLC, **(iv)** at the **RLC/RLC** interface that involves direct asymmetric apolar interactions between helices 1 and 2 of the BH RLC and the N-term extension (M20, F21) and loop2 of the FH RLC (**Fig. 2e**); **(v)** at the **RLC/cc** interface where HC residues F834-E846 adopt drastically different positions (**Extended data Fig. 7c**). While all these _BH_HC residues mediate interactions within the _BH_RLC, only the first part of the _FH_HC residues interact within the _FH_RLC via different contacts than those formed in the BH head. After Pro838, _FH_HC residues form stabilization contacts at the surface of the _BH_RLC, in particular via L839 and R845 (**Extended data Fig. 7c**).

Interestingly, all these interfaces in cardiac IHM are quite modest in surface area and involve several electrostatic interactions, as shown by the electrostatic profile of the footprint (**Extended data Fig. 8**). Their formation at different places of the IHM maintains the two heads together, yet their relative position is distant which allow variability and leads to lower resolution for the lever arm region compared to the motor domains of the IHM. In addition, most of the contacts made are dynamic in nature making it difficult to visualize side-chains at the interface. As previously described for the Converter/ELC interface^21^, these electrostatic interactions are labile and can arrange differently. The structure demonstrates the generality of the concept of “musical chairs”^21^ as it is found at different interfaces and hinges of flexibility of the cardiac IHM. Finally, the interfaces formed in the IHM include key-elements for actin interaction and mechano-chemical transduction (Transducer and Converter). While interactions with S2 and constraints between the lever arms efficiently result in positioning the heads away from the actin filament, blocking the dynamics of regions involved in force production also contribute to the efficient shutdown of myosin activity.

## Cardiac and Smooth IHM differ

Surprisingly, at high resolution, the β-cardiac IHM and the SmMyo2 IHM strongly diverge in structure. This was totally unexpected since from the low and medium resolution structures, the IHM was thought to be conserved in all class-2 myosins (reviewed by^31^).

The first major difference concerns the orientation of the heads within the motif: when the cardiac and the SmMyo2 IHM are aligned on the BH, the cardiac FH is rotated ∼20° anticlockwise compared to the smooth FH (**Fig. 3a**). At the head/head interface, the conformations of the FH β-bulge and HO-linker differ and the interaction they mediate are distinct among these two IHMs (**Fig. 3b**, **Extended data Table 2**). In addition, the _FH_Top-loop adopts different conformations in Smooth and Cardiac FH heads and distinct interactions are formed by the _FH_Converter with the similar _BH_PHHIS region of the U50 subdomain in the two IHMs (**Fig. 3b**). In fact, the sequence alignment of elements involved in the interfaces shows major differences among Myo2s (**Fig. 3c**), which are linked to changes in structure. For example, the Top-loop of SmMyo2 is one residue shorter and comprises **(i)** a charge reversal at the position E732 (replaced by a lysine in SmMyo2); **(ii)** replacement of the key I736 by a methionine (**Fig. 3c**). Such differences prevent conservation of interactions. Major sequence differences in regions involved in the head/head interfaces are also found in the HO-linker and the β-bulge. Interestingly, from sequence comparison, we can anticipate that the IHM of skeletal muscle Myo2 (SkMyo2) is close to that of cardiac myosin while that of non-muscle myosin 2a (NM2a) would be similar to the SmMyo2 IHM.

**Figure 3.**
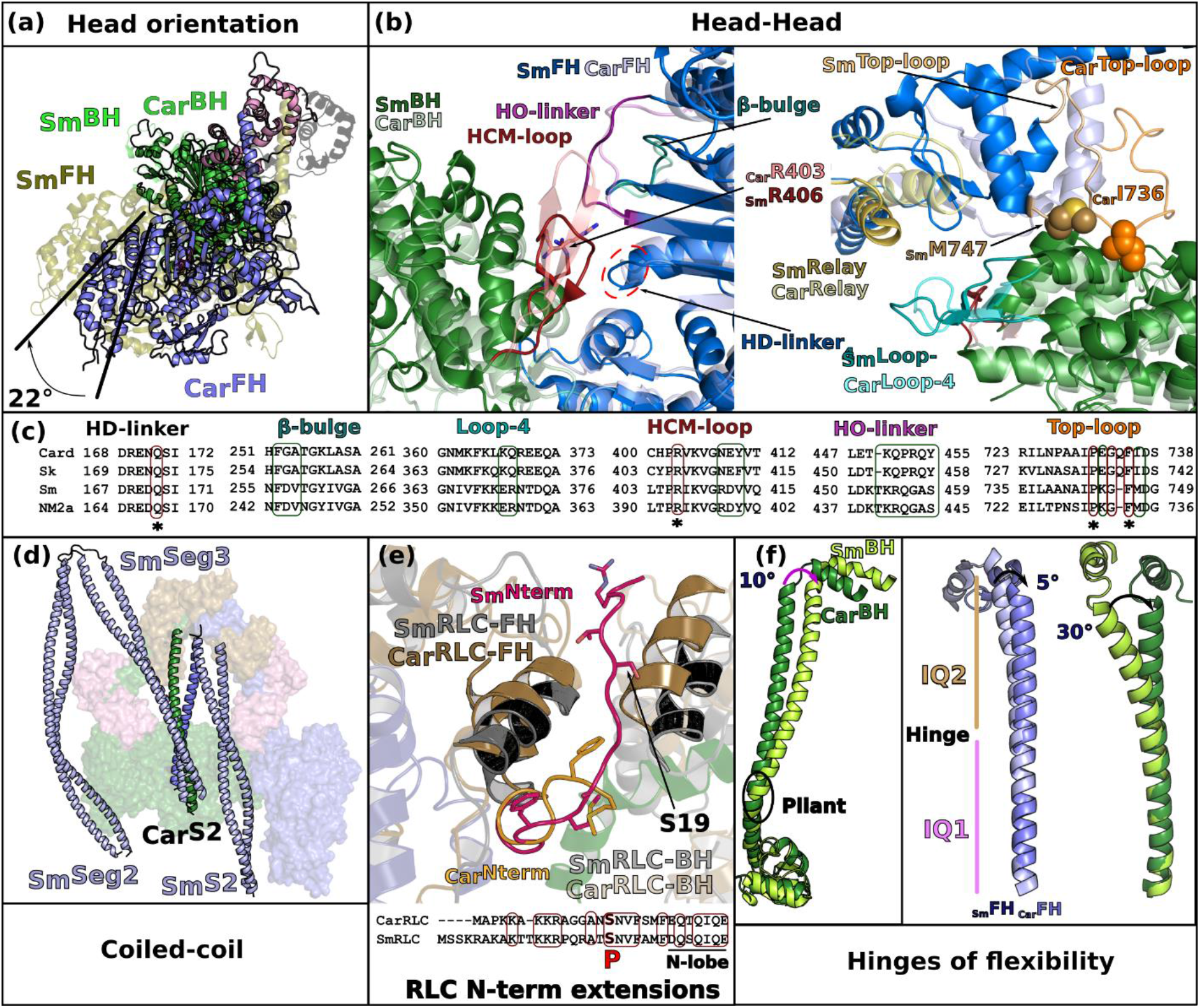
– The sequestered state of β-cardiac myosin and smooth muscle myosin (SmMyo2) differ greatly. **(a)** Relative orientation of the BH and FH heads in Smooth (Sm, PDB code 7MF3, ^15^) and Cardiac (Card) IHMs. SmMyo2 and cardiac myosins are aligned on the BH (green), the FH of cardiac myosin is rotated 22° anticlockwise compared to the FH of SmMyo2. **(b)** Comparison of the head-head interfaces in SmMyo2 and cardiac myosin, two views are represented. On the left, aligned on the FH motor domain, on the right, aligned on the BH motor domain. The cardiac IHM is represented in transparency. Note the difference in the HCM-loop on the left, and on the _FH_Relay (yellow) and _FH_Converter positions (Blue) on the right. **(c)** Sequence alignment of the different connectors and elements involved in the interactions showing the divergence: Card_Hs (Homo sapiens β-cardiac myosin); Sm_Gg (Gallus gallus smooth muscle myosin); Sk_Hs (Homo sapiens fast skeletal muscle myosin); NM2a_Hs (Homo sapiens nonmuscle myosin 2a). Regions of interactions are in colored box: conserved in red, divergent in green. **(d)** Comparison of the position of the coiled-coils on the surface of the heads between cardiac myosin and SmMyo2. **(e)** Comparison of the RLC-RLC interface, the models are aligned on the IQ2 of the FH. A sequence alignment of the N-term extension of the RLCs is shown with conserved regions between smooth muscle RLC (SmRLC) and cardiac RLC (CarRLC) boxed in red. The phosphorylation is represented in red with “P”. **(f)** Comparison of the angles at the hinges of flexibility: pliant region, aligned on the Converter (on the left), FH and BH lever arm, aligned on IQ1 (on the right).

The second major difference is the positions of the S2 coiled-coil (S2). In the SmMyo2 IHM structure, S2 is located close to the FH head and the IHM is strengthened by interaction with two additional coiled-coil segments (Seg2 and Seg3). No such segments stabilize β-cardiac myosin, rendering the IHM motif less stable when isolated from filaments^13–15^. Surprisingly, the cardiac S2 interacts extensively with the Mesa of the BH, in drastic contrast to the position S2 adopts in the SmMyo2 IHM, which only allows for a few interactions with the _FH_Loop2. In fact, it is Seg3 that interacts with the BH Mesa in SmMyo2 IHM (**Fig. 3d**).

More importantly, the detailed comparison of the RLC/RLC interface reveals the structural differences underlying the distinct modes of regulation of the cardiac and SmMyo2 IHM formation. Both _Car_RLC and _Sm_RLC are phosphorylatable in a charged and disordered N-terminal extension (N-term extension) which is not conserved in sequence (**Fig. 3e**), although from the globular N-terminal lobe of the RLC, the position of the two phosphorylatable serines S19 in _Sm_RLC and S15 in _Car_RLC is conserved (**Fig. 3e**). In SmMyo2, RLC phosphorylation acts as an on/off switch to disrupt the SRX and activate the muscle^32^. This was explained in the Smooth IHM structures^13,15^where the _BH_RLC N-term extension interacts with seg3 and the _FH_RLC N-term extension is directly part of the RLC/RLC interface. S19 Phosphorylation thus disrupts these IHM interfaces, switching on the motor (**Fig. 3e**). In cardiac myosin, RLC phosphorylation is not strictly necessary to activate cardiac muscle but modulates its activity^33^). The cardiac IHM structure explains this more moderate effect on IHM interfaces, beyond the fact that seg3 is not part of stabilizing the cardiac IHM. The RLC/RLC interface is drastically different in the cardiac IHM: the _BH_RLC extension is not part of it and the _FH_RLC N-term extension is only found at the periphery of this interface (**Fig. 3e**). No density allows precise positioning of the _FH_RLC S15 as it is not a major part of the RLC/RLC interactions. Thus, phosphorylation would only modulate the stability of the IHM in cardiac. This major structural difference in how the smooth and cardiac IHM form underlies the distinct regulation in striated and smooth muscles.

Finally, the differences in the hinges of flexibility of cardiac and SmMyo2 myosins are required to allow the distinct but precise asymmetric orientation of the heads in the two IHMs. The flexibility and the kink at the pliant region are much larger in the BH head compared to the FH, which adopts a relatively straight lever arm. However, the BH pliant region is 10° more kinked in SmMyo2 compared to cardiac (**Fig. 3f**). The kinks of the HC at the IQ1/IQ2 region are different in the two IHMs. There is an angular difference of 5° between smooth and cardiac FH heads and 30° between the BH heads respectively (**Fig. 3f**). The ELC/RLC interactions are thus different both for the BH and FH heads. This validates that these regions are specific ankles of flexibility required for the formation of the IHM (**Fig. 3f**).

## Consequences on muscle physiology

In the thick filament, the IHM motifs are organized in crowns with different symmetries depending on the muscle and organism considered^12^. This different organization implies distinct inter-crown interactions that could not be resolved precisely with low-resolution models of the relaxed thick filaments^31, 34^ The 28 Å EM structure of the human cardiac filament revealed repeats of three crowns, crown 2 being less ordered^10^. Regarding the tilt or orientation of each crown, crown 1 is rotated 19° in a clockwise direction as viewed from outside the filament compared to crowns 3 and 2, which are similar^10^ (**Fig. 4a**). This allows a strong interface to be formed between crowns 1 and 3, but not between crowns 3 and 2. The distance between the crown 2 and crown 1 heads is too large for interactions to be formed, thus resulting in a perturbation of the helical symmetry^35–37^.

**Figure 4.**
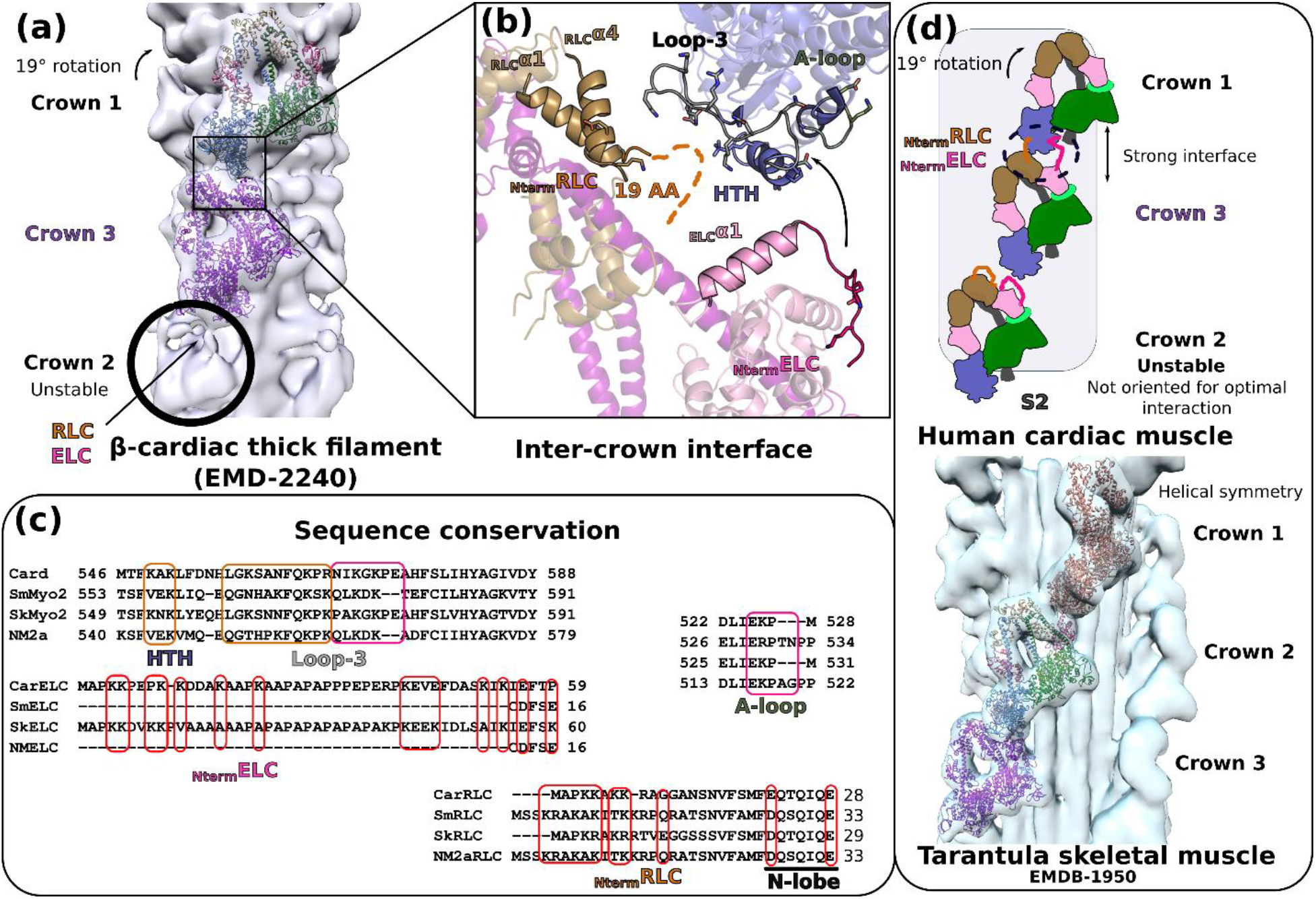
– Organization of the thick filament. **(a)** Fit of the IHM motifs in crowns 1 and 3 using the 28 Å resolution map of the relaxed human cardiac thick filament (EMD-2240^15^). Crown 2 is less stabilized and less defined in density. A tilt of 19° is observed for crown 1 compared to crowns 2 and 3. **(b)** Zoom in the interface between crown 1 and 3 that was previously thought to involve mainly the RLC of crown 3. Our model indicates that the ELC N-lobe and the N-terminal extensions (Nterm extensions) of both the ELC and the RLC of crown 3 could interact with the L50 HTH and loop 3 of crown 1. The 49 amino-acid Nterm extension of the ELC can reach the activation loop (A-loop) and the helix-turn-helix (HTH). **(c)** Analysis of the sequence conservation of the regions involved in the inter-crown interface. **(d)** Schematic overview of the differences stabilizing the sequestered states in striated and smooth muscle. In cardiac muscle (upper panel), the inter-crown interface stabilizes the IHM of crowns 1 and 3, while crown 2 is less stabilized and less defined in the density because (i) it is not oriented to establish this interface and (ii) the crown 2/crown 3 inter-crown distance is longer compared to crown 3/crown 1. A fit of the IHM cardiac model in the purely helical relaxed cardiac skeletal muscle^38^. The helical symmetry induces different inter-crown interfaces compared to cardiac relaxed filament.

We placed the high-resolution structure of the cardiac IHM in the thick filament electron density designated for crown 1 and 3^10^, which allowed us to describe the precise structural elements involved in the most stable inter-crown interface (**Fig. 4a**). Interestingly, crown 1 and crown 3 heads can interact via three actin-binding elements of the FH head of crown 1, (i.e. the helix-turn-helix (HTH), activation loop (A-loop) and loop-3), and the N-terminal extensions of the _BH_RLC and the _BH_ELC of crown 3 (**Fig. 4b**). These extended regions contain several charged residues that can mediate long-range interactions and concur in stabilization of the two crowns in their respective orientation (**Fig. 4b**). Sequence alignment of elements of the interface confirm a specific conservation between striated muscle myosins (cardiac and skeletal) that is not found in smooth muscle and non-muscle Myo2s (**Fig. 4c**). The ELC that binds SmMyo2 and NM2a have no N-terminal extension and their RLCs have longer and divergent N-term extensions compared to cardiac RLC (**Fig. 4c**). As was previously suggested from a low resolution model of the relaxed muscle thick filament^10^, our model shows that the RLC is part of the Crown1/Crown3 interface, providing now in addition a potential role for the N-terminal lobe of cardiac ELC for stabilizing these IHMs. The BH RLC S15 phosphorylatable residue would also be found close to the inter-crown interface, suggesting how phosphorylation of the crown 3 BH RLC head might weaken the cardiac IHM stability and modulate the number of active myosin heads. In other muscles with different symmetry for the thick filament (e.g. tarantula skeletal muscle which harbors a perfect helical symmetry^38^, the inter-crown interface differs and does not involve the RLC (**Fig. 4d**; **Extended data Fig. 9**). Thick filament geometry may thus induce different types of inter-crown regulations.

The N-terminal extension of _Car_ELC is also involved in direct interaction with actin and increases cross-bridge kinetics (reviewed by ^39^). Sequestering it in the inter-crown interface would be an additional mechanism to block interaction with actin. Interestingly, most of the elements involved in actin binding are part of either the intra-molecular interactions within an IHM or the inter-crown interface. The inter-crown stabilization interactions challenges the concept of a free-head^8^ and the idea that the IHM of cardiac myosin is less stable than that of SmMyo2. In the cardiac thick filament, indeed stabilizing partners such as MyBP-C or Titin^10^ could further stabilize the heads and the inter-crown interactions. Thus, as many as half of the cardiac heads may be shut down in the off state even during systolic contraction of the heart^6, 10^. In contrast, strain upon contraction could disrupt these thick filaments stabilization interactions and would increase the number of heads recruited for optimal heart contraction.

## Inherited cardiomyopathies and therapies

One of the major predictions to explain the hyper-contractility of the cardiomyocytes during the early stages of hypertrophic cardiomyopathies (HCM) is the increase in the number of heads available for contraction^18^, and specifically due to the destabilization of the IHM by point mutations responsible for the disease^21, 27, 28, 40^. Exploiting structural knowledge of the myosin allostery promoting force production and low-resolution models of the IHM, HCM mutations have been grouped in different classes^27^ to predict how mutations may influence contractility. Consistent with this hypothesis, many HCM mutations studied with purified human β-cardiac myosin constructs have indeed been shown to directly decrease the SRX and diminish the stability of the IHM^17, 28, 41,42^Differences in the number of heads available to interact with actin are thus likely to be a major cause of the disease, although each mutation can also affect differently the intrinsic force produced, the ATPase rate, and/or the velocity of movement of actin over myosin in the *in vitro* motility assay^17, 20, 21, 28, 41, 42^.

The cardiac IHM structure shows that a number of mutations indeed lie close to the IHM stabilizing interfaces (**Extended data Fig. 10**). With this long awaited structure, the prediction for mutations in critical regions such as the Converter can be clarified (**Extended data 1**), and a more accurate prediction of the effects of HCM and DCM mutations can be released (**Extended data 1**). This high-resolution structure now provides a precise distinction between **residues directly involved in the intra-molecular IHM interactions** (**Extended data Table 2, Extended data 1**) and those found nearby that affect these interfaces indirectly via their influence on the conformation of critical structural elements required for the stabilization of the IHM. The high-resolution IHM structure provides also details on how mutations in the light chains or at the main interfaces can disrupt the IHM, that could not previously have been interpreted. Previous low-resolution models used for mutation prediction^42^were all unable to model correctly the **hinge of flexibilities and light chains** and how mutations could affect the **inter-crown interface**. An example here is _ELC_A57G, a severe infantile mutation **in the ELC light chain**^43^ that leads to a decrease of the SRX in mouse cardiac fibers^44^. The IHM structure indicates how the replacement of A57 to a glycine would destabilize helix _ELC_α1 and _ELC_Loop-1, and thus the ELC/RLC interface. In fact, these ELC structural elements are crucial for stabilizing distinctly the hinges of the two lever arms of the IHM. In addition, this _ELC_A57G mutation can directly affect the inter-crown interactions (**Extended data 1,** **Fig. 5a**). Our analysis also indicates that the inter-crown and the RLC/RLC interfaces will also be weakened by HCM mutations **in the RLC N-term extension**, such as A13T, F18L and E22K (reviewed by ^43^) (**Extended data 1;** **Fig. 5a**). Mutations in the **actin binding elements of the L50** (Loop3, activation-loop (A-loop) or helix-turn-helix (HTH)), initially interpreted as disrupting motor function, are in fact in the <u>inter-crown interface</u> and could thus also dysregulate the number of heads participating in contraction (**Fig. 5a**). Interestingly, while six HCM mutations (two located in the A-loop, two in loop-3) are predicted to alter the inter-crown interface, this region is enriched with five DCM mutations (**Fig. 5a**). The consideration of the inter-crown interface leads to the definition of a new class of cardiomyopathy causing mutations, which mainly affect the IHM stability by its stabilization on the filament level via interactions between IHM crowns or with thick filament associated proteins such as MyBP-C and titin. Higher resolution structures of the thick filament will be required to further analyze the effect of such mutations.

**Figure 5.**
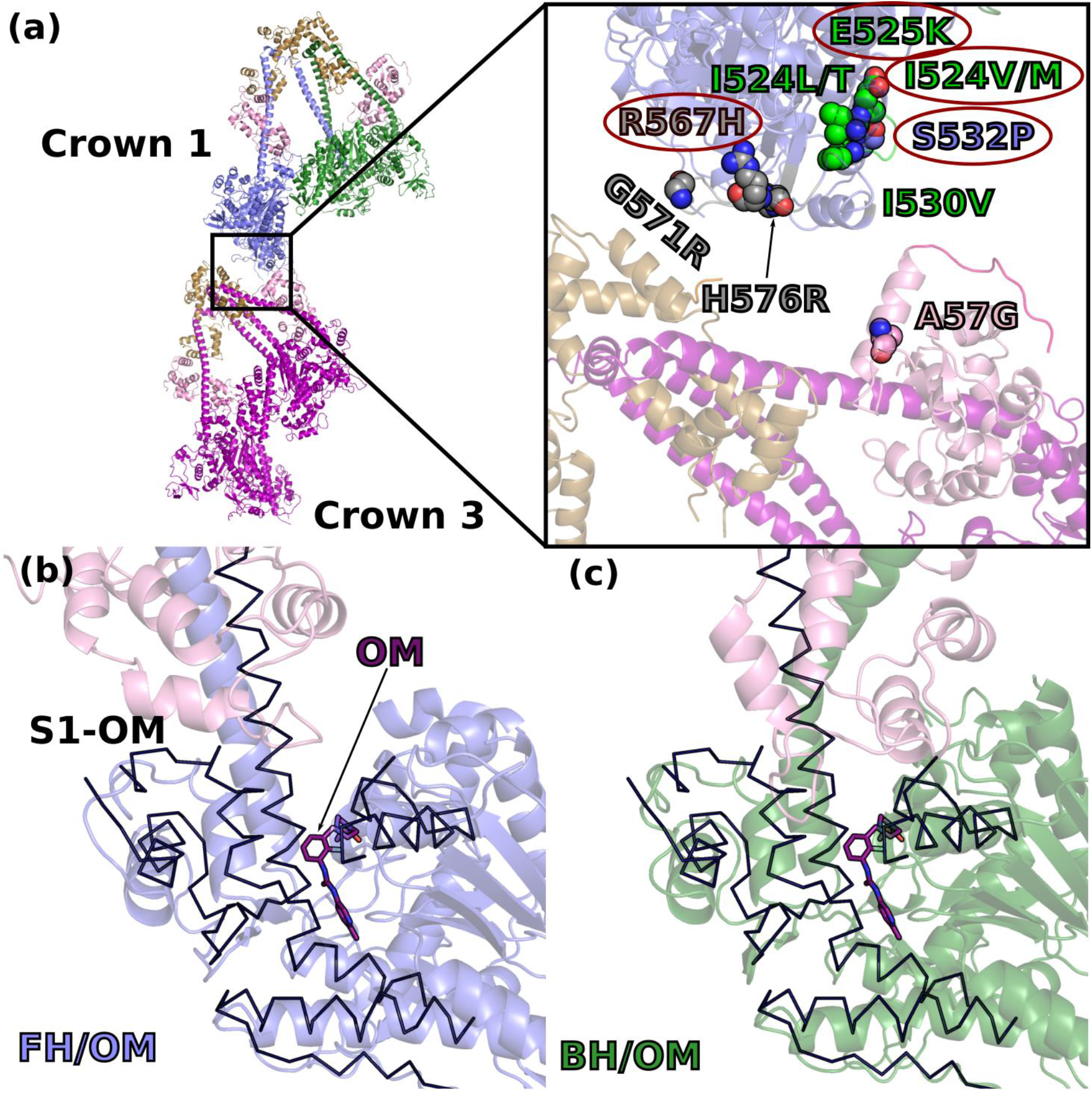
– Molecular determinisms underlying hypertrophic cardiomyopathy (HCM) and dilated cardiomyopathies (DCM). **(a)** DCM (circled in red) and HCM causing mutations predicted to destabilize the inter-crown interface or the IHM are represented on a cartoon representation of crown 1 and 3. Zoom at the inter-crown. **(b)** and **(c)** show a superimposition of the structure of the S1 fragment of cardiac myosin bound to the activator Omecamtiv mecarbil^24^ (PDB code 5N69, PPS-S1-OM) superimposed on the motor domain of each head FH and BH respectively. PPS-S1-OM is represented as a black ribbon and shows how OM induces an orientation of the lever arm that is not compatible with the requested orientations for the FH or the BH when the IHM is formed.

Small molecules modulating cardiac myosin force production are among the most promising therapies for inherited cardiac diseases. The activator Omecamtiv mecarbil (OM) was reported to destabilize the IHM^22^. The OM-bound cardiac myosin structure ^24^ showed that OM binds in a pocket of the cardiac pre-powerstroke state located between the motor domain and the Converter. The closure of this pocket upon OM binding introduces a hinge that shifts the Converter orientation prior to the pliant region. This now makes these two OM-bound heads incompatible with the sequestered state (**Fig. 5b, 5c**), explaining the reduction in the IHM formation in the presence of this activator. The IHM structure will be essential to identify different classes of inhibitors and whether they are compatible or not with the formation of the IHM depending on which pockets they bind to.

## Conclusions

The high-resolution structure of the sequestered state of β-cardiac myosin is paramount for understanding thick filament regulation. The unexpected divergence with the SmMyo2 IHM structure reveals that while myosin regulation is conserved in evolution, distinct sequences have evolved from a common ancestor to fulfill the requirements of distinct fiber types and muscle functions (**Fig. 4d**). Interestingly, most of the interfaces involve actin-binding elements that are variable and these differences were reported to tune **(i)** myosin force production and **(ii)** the orientation on the actin tracks^1, 45, 46^. It is tempting to speculate that actin-binding elements are part of a super-regulation mechanism, also regulating the sequestered state stability. Future high-resolution studies of other IHMs will provide insights to decipher how the divergence of the IHM motif evolved to also fit different types of Ca^2+^ or phosphorylation regulation, together with fulfilling different types of muscle contraction properties.

The IHM structure provides the missing puzzle piece to understand the molecular mechanism of inherited cardiomyopathies. As for β-cardiac myosin, MyBP-C is also a hotspot of HCM mutations^47^. It was reported to increase the SRX through direct interactions with cardiac myosin^28, 48^. MyBP-C and Titin are positioned to influence the myosin crowns in the relaxed thick filament^10^. Future structural studies with these partners will further complete the modelling of how mutations can alter SRX regulation. With a near-atomic model of the IHM, the mode of action of future treatments of cardiomyopathies can also be deciphered more precisely. The availability of the cardiac IHM structure will pave the road towards the rational design of novel therapeutics against distinct families of inherited muscle disease.

## Acknowledgements

This work was funded by grants NIH RM1GM131981-01 (J.A.S and A.H.), ANR-21-CE11-0022-01 (A.H.), and NIH R01GM33289 (J.A.S.). NN acknowledges postdoctoral funding from AHA (908934) and Stanford MCHRI (<u>1220552-140-DHPEU</u>). DP acknowledges postdoctoral funding from AHA.

## Authors contributions

Conceptualization and design of the research: JRP, AH, JAS and KMR. Protein expression, purification and characterization: NN, DP. Negative staining: AG, CK, JRP. Set up and conditions screening to prepare the CryoEM grids: AG, JRP. Technical support, DM. Data collection and processing: AG under the supervision of EK. Model building and refinement: JRP, DA, AH. Model analysis: JRP, DA, AH. HCM mutation analysis and update of the table: JRP, AH, KMR, JAS. Manuscript writing JRP, AH with the help of the other authors. Project administration: JRP, AH, JAS and KMR.

## Data availability

The atomic model is available in the PDB ^49^ under the code 8ACT. The cryo-EM maps Map 1 and Map 2 are available in the EMDB database^50^ under the accession numbers EMD-15353 and EMD-15354 respectively.

## Materials and methods

### 15-hep production and purification

The human β-cardiac myosin construct 15-hep HMM was produced using a modified AdEasy™ Vector System (Qbiogene, Inc, Carlsbad, California, USA). The 15-hep cDNA consists of residue 1 to 942 of *MYH7*, encompassing S1 and the first 15 heptad repeats of the proximal S2 coiled-coil region. This sequence is followed by a GCN4 leucine zipper to ensure dimerization, a flexible GSG linker, and finally a carboxy-terminal PDZ-binding octa-peptide (RGSIDTWV). The adenoviral vectors expressing the heavy chain myosin and the human ventricular essential light chain (ELC) (*MYH3*) containing a TEV protease cleavable N-terminal FLAG tag (DYKDDDDK) were co-expressed in differentiated mouse myoblast C2C12 cells (purchased from ATCC). C2C12 cells were infected with the adenoviral vectors 48-60 hours after differentiation and harvested 4 days post infection in a lysis buffer with the following composition: 20 mM imidazole at pH 7.5 containing 50 mM NaCl, 20 mM MgCl_2_, 1 mM EDTA, 1 mM EGTA, 10% sucrose, 1 mM DTT, 3 mM ATP, 1 mM PMSF and Roche protease inhibitors. Harvested cells were immediately flash frozen in liquid nitrogen and stored at -80°C.

For purification, frozen cell pellets were thawed at room temperature, and supplemented with 3 mM ATP, 1 mM DTT, 1 mM PMSF, 0.5% Tween-20 and Roche protease inhibitors before lysis with 50 strokes of a dounce homogenizer on ice. The lysate was incubated on ice for 20-30 mins to allow for the assembly of native mouse full length myosin into filaments before clarification; filament formation is promoted by the high MgCl_2_ and low salt content of the lysis buffer. The lysate was clarified by spinning at 30,000 RPM in a Ti-60 fixed-angle ultracentrifuge rotor for 30 minutes at 4°C. All subsequent steps were carried out in a cold room set at 4°C. The supernatant was incubated on a nutating rocker with anti-FLAG resin for 1-2 hours, followed by low-speed centrifugation to settle down the protein-bound resin. The lysate was decanted, and protein-bound resin was washed with >10-fold excess of a wash buffer (20 mM imidazole at pH 7.5, 5 mM MgCl2, 150 mM NaCl, 10% sucrose, 1 mM EDTA, 1 mM EGTA, 1 mM DTT, 3 mM ATP, 1 mM PMSF and Roche protease inhibitors). Native mouse regulatory light chain (RLC) was then depleted by incubating the resin with RLC depletion buffer (20 mM Tris-HCl at pH 7.5, 200 mM KCl, 5 mM CDTA pH 8.0, 0.5% Triton-X-100 supplemented with 1 mM ATP, 1 mM PMSF, and Roche protease inhibitors) for 75 mins on a nutating rocker. The resin was washed with wash buffer and incubated with human RLC, purified from *E. coli* as previously described^4^, for ∼3 hours. Finally, unbound human RLC was washed away, and the resin was nutated overnight in wash buffer containing TEV protease to cleave the hexameric myosin assembly from the resin. The next morning, the protein was further purified using anion-exchange chromatography. The myosin in solution was separated from the stripped-off resin using a Micro Bio-Spin™ Chromatography Column (Bio-Rad) and loaded on a 1 mL HiTrap Q HP column (Cytiva) attached to an FPLC. The protein was eluted with a gradient of 0-600 mM NaCl (in a buffer containing 10 mM imidazole at pH 7.5, 4 mM MgCl_2_, 10% sucrose, 1 mM DTT and 2 mM ATP) over 20 column volumes. The fractions corresponding to the protein peak were run on a 12.5% SDS-PAGE and pure fractions (devoid of the contaminating full length mouse myosin) were pooled. The protein was concentrated to ∼10 mg/ml using Amicon Ultra centrifugal filter units, and flash frozen as 20-40 uL aliquots.

Single ATP turnover experiments from Jim Spudich lab (manuscript in preparation) and others^7^ confirm that 15 heptads of the proximal S2 tail allow myosin to adopt the folded-back SRX state in solution.

#### Cryo-EM data collection and processing

The sample, was diluted at 0.3 mg/ml concentration in 10 mM imidazole pH 7.5, 2 mM MgCl_2_. 0.5 mM Mg.ATP was added and the mix was incubated 15 minutes at room temperature. 3 µl was applied to glow discharged UltrAuFoil 1.2/1.3 holey gold grid and vitrified in a Mark IV Vitrobot (ThermoFisher). Grids were imaged in ESRF’s CM01 facility using a Titan Krios microscope (ThermoFisher) at 300 keV ^8^ with a K3 direct electron camera at 0.84 Å per pixel. 5154 movies were collected with 40 frames each and a dose of 0.98 e-/Å^2^ per frame. After beam-induced motion correction with Motioncor2^9^ and Contrast transfer function (CTF) estimation with Gctf^10^, 4423 micrographs were selected for further particle analysis performed in cryoSPARC^11^. A total of 493179 particles were selected after automatic picking and the first round of 2D classifications. These particles were used to generate several *ab initio* reconstructed 3D models that were subsequentially used for a 3D heterogeneous refinement. At the end of this first round of refinement, 213596 particles were unambiguously attributed to the sequestered state. These particles were further processed with 3D refinement, using the cryoSPARC homogeneous refinement algorithm^11^, giving final global resolutions based on the gold-standard Fourier shell correlation (FSC = 0.143) criterion^12^ of 3.6 Å (map 1). To further improve the resolution of the heads region, a homogeneous refinement using a mask covering the heads region alone was carried out. This final refinement produced a 3.2 Å resolution map of this region (map 2). Local resolutions of the density maps were calculated in cryoSPARC^11^.

#### Model building and refinement

The head/Converter/ELC region was rebuilt based on the bovine β-cardiac myosin PPS complexed to Omecamtiv mecarbil^2^ (PDB code 6N69) without solvent or ligand. The coiled-coil S2 region was fitted from the crystal structure^13^ (PDB code 2FXM). The IQ2/RLC region was obtained by homology modeling based on the SmMyo2 IHM^1^ (PDB code 7MF3). Each part was fitted in the density map and the model was built and optimized with the module Quick MD Simulator from CHARMM-GUI^14, 15^. Steps of real space refinement were performed against map 1 with Phenix^16^. During the procedure, the head region (residues 1-781) which was at higher resolution was subjected to reciprocal space refinement against map 2 with Refmac^17^. An independent run of molecular dynamics was performed in order to improve the statistics at the interfaces and to compute realistic interactions. The system was built with the CHARMM-GUI/Quick MD simulator module^14, 15^. Only the interfaces (head/head, coiled-coil/head) of the model were relaxed in a box containing explicit water (TIP3P) and salt (150 mM KCl) in the CHARMM36m^18^ force field. 10 ns of simulation was performed in GROMACS (version 2018.3)^19^. A final run of real space refinement was performed against map 1. Quality indicators were calculated with Phenix, including Molprobity^20^ and EMRinger^21^.

#### Interface analysis

The analysis of the interfaces was performed automatically with PDBsum^22^. Interactions were manually checked and extended up to 4.5 Å to discuss long range weak interactions. We chose to discuss the realistic interfaces of the IHM because we computed it with molecular dynamics. While some ambiguity exists at times in these interfaces, the electron density is good for the main chain and the large hydrophobic side-chains. As discussed in the text, most of the interfaces of cardiac IHM contain however labile side-chains (musical chairs) that can adopt more than one conformation. Lower EM density for such side chains are then apparent, even in the high-resolution regions such as the head-head regions.

## Extended data 1

With a detailed analysis of the high-resolution structure of the β-cardiac IHM, we can provide here a detailed evaluation of the effects of cardiomyopathy mutations. The precise analysis of these mutations on the interfaces was impossible in the absence of a high-resolution structure for the IHM. The fact that mutations can be located in connectors such as the Transducer or the Converter also complicate their analysis as they have an impact in motor function^46^. With the IHM structure, the impact on the stabilization of the heads can be precisely assessed as we have access to the conformation these flexible elements adopt to form the IHM interfaces. With a detailed analysis of a few HCM mutations and the structural impact for the IHM and for the function of force generation, we can now more clearly distinguish the different classes of mutations that can lead to the cardiomyopathies.

## Mutations in the heads

In the last years, several mutations located in the motor domain have been analyzed in the Spudich laboratory and in other groups. Some mutations were directly reported to decrease the SRX state: R249Q; H251N; D382Y; R403Q; V606M; P710R and R719W^4, 23, 34, 47, 48^. The IHM structure now provides detailed structural insights to identify how mutations can lead to the development of the HCM phenotype.

### Some of these mutations are located at the interfaces and would thus directly impact the IHM stabilization

**R403Q** is a hotspot of mutations^46, 49^, which would directly affect the head-head interface by destabilizing contacts between the _BH_HCM-loop, the _FH_HO- and the _FH_HD-linkers (see main text and **Figure 2b**; **Extended data Table 2**). This explains the significant decrease of the SRX measured with this mutant^23, 47^. Even if R403 is not directly involved in the actomyosin interface in Rigor^50^, the alteration of the local structure and dynamics of the loop is responsible for the 2-fold reduced binding affinity of the mutant for the actin track^51^.

**D382Y** is not located on the Mesa but on the so-called _BH_PHHIS surface of the _BH_U50 subdomain. This surface residue is directly involved in polar interactions with the _FH_HD-linker (**Figure 2b**; **Extended data Table 2**) and we predict that the mutation would have little effect on the structure of the head itself. The fact that the IHM structure predicts that the mutation would destabilize the head-head interface is consistent with data indicating a 46% reduction of the SRX^34^, with only modest alteration of the mechanochemical properties of the motor carrying the mutations^52^.

**R453C** (HO-linker) is also a direct part of the interface between the _BH_motor domain and the S2 coiled-coil (**Figure 2c**; **Extended data Fig. 10a**). Interestingly, this mutation would greatly affect a second interface of the IHM when present in the FH head: the interactions between the HO-linker of the Mesa of the FH and the _BH_HCM-loop (**Extended data Table 2**). While the effect of the mutation on the SRX has not been measured yet, it decreases the affinity of the S1 fragment for the S2 coiled-coil^4^, consistent with our IHM structure. Since this mutation is located in a linker of the Transducer, it is predicted to alter the structural transitions between the states and thus it is not surprising that it changes motor properties^29^.

### Some of the mutations are not located at the interface but close enough to induce indirect effects and destabilize the IHM

**H251N** and **R249Q**, two mutations of the β-bulge of the Transducer, lead to a 41% and 59% decrease of the SRX as well as a decrease in the affinity of the S1 head for the S2 coiled-coil ^4, 34^. Given the essentiality of the Transducer in conformational changes during the motor-cycle, the role of these mutations in force generation has been demonstrated for H251N^53^ and is also likely for R249Q^54^ although this has not yet been studied in human. The R249 and H251 side chains are in the vicinity but are not involved in direct interactions stabilizing the IHM (**Extended data Fig. 10**). In the FH head, this Transducer region interacts with the BH-HCM loop (near _BH-HCM-loop_E409). In the BH head, the Transducer interacts with the S2 coiled-coil. Destabilization of the IHM may result from local changes in the dynamics of main chain or side chain of the β-bulge, which would then impact the contacts made in the IHM and its stability on two interfaces that involve the Transducer: _BH-_Tranducer with the S2 coiled-coil and _FH-_Transducer with the HCM-loop. Since H251 and R249 are involved in polar interactions between strands of the Transducer, the presence of the Asn or a Gln side chain instead of H251 or R249 may also alter the stability of the pre-powerstroke state which would indirectly affect the formation and stability of the IHM, as previously proposed^46^. Thus, the structure of the IHM provides much more detailed hypotheses of the role of the mutations than previously possible, which can guide further investigations.

**R663H,** located at the end of the HW helix on the surface of the myosin head, is close to the Mesas of the FH and the BH. Its surface location would predict that a mutation at this position should not affect the mechano-chemical parameters of the motor. Kinetics experiments demonstrate that this mutation has indeed no effect on the fundamental parameters of the myosin motor, while a major reduction in the SRX is found^23^. The BH R663 side chain in the IHM is close to the _BH_head/coiled-coil (_S2_D896/D900) interface while the FH R663 side chain is close to the _FH_head/loop4 (_BH-loop4_E370) interface, but the current cryoEM map density does not support direct interaction. The charge-removal introduced by the mutation would change the global charge of both Mesa and thus likely remotely affect the IHM formation, possibly also by affecting how this residue interacts with other regulatory proteins of the thick filament.

**R719W** is located in the Converter, in close vicinity of the Top-loop, a key-element in the head/head interface. While previously debated among IHM models, the IHM structure clearly defines how the FH-Converter is positioned to stabilize interactions with the BH head. This mutation is predicted to destabilize the Top-loop and alter its dynamics and conformation, thus altering the _BH_PHHIS/_FH_Converter interface, resulting in a 68% decrease of SRX^34, 47^. Interestingly, this mutant only has modest effects on kinetics parameters of the S1^52^ while a 20% decrease of intrinsic force was reported^55^. This is consistent with the location of the mutation that would not disrupt the Converter swing but would alter the Converter/ELC interface and thus the flexibility of the lever arm^3^ (Robert-Paganin *et al*., 2018).

**R723G and G741R** are also close to the Top-loop and would have similar outcomes to R719W. This is confirmed by biochemical assays demonstrating that these mutants have only small effect on S1 mechanochemical properties^55^. Thus, the effects of these mutations are likely only on the _BH_PHHIS/_FH_Converter interface of the IHM by altering the conformation of the Top-Loop, as previously proposed via molecular dynamics predictions of the Converter^46^. However, the quantitative effect of such mutations on the SRX are not yet available.

### Finally, a group of mutations affecting the SRX are located far from the interfaces but in regions important for the allosteric conformational changes of the motor upon force production

**P710R** is located in the core junction of the Converter, far from all interfaces that stabilize the IHM. However, it can greatly alter the conformation of the SH1-helix leading to drastic alterations of the allostery within the motor. In addition, it can also alter the stability or the conformation of the PPS state that both heads must adopt to form a stable IHM. We predict that in this case, the alteration of the PPS state could also lead to a change in the lever arm position that could result in difficulty in forming the IHM conformation. Thus, the mutation leads to both a decrease of the SRX and a dramatic alteration of motor functions (**Extended data Fig. 10a**)^48^.

**V606M** is also far from the interface but located in the U50 core close to the strut and the actin binding cleft and also close to Loop2, an element essential in the first step of actin binding (**Extended data Fig. 10a**) ^56^. The replacement of a Val by the bulkier Met may alter the structural rearrangements during the powerstroke and the actin binding cleft opening and closure. This may induce severe consequences on mechano-chemical properties of the motor such as association to actin and force output. The V606M also decreases the SRX ^47^ and this effect is likely due to both a potential alteration of the PPS state but also on the nearby _BH_PHHIS surface.

As proposed previously, the mutations in the heads can have effects on IHM stability and/or on the ability of the motor to produce force^46^. These two parameters should be taken into account when discussing the molecular consequence of inherited cardiomyopathy mutations. Interestingly, some of the mutations analyzed here are located in the Transducer or the Converter, central elements in force production and allosteric communication of the motor. HCM mutations would thus have two ways of destabilizing the IHM: **(i)** direct effect on the interfaces or on the conformation of elements of the interfaces and **(ii)** remote effect on the stability of the PPS. Since the formation of the IHM requires the two heads in PPS, case (ii) alters the ability of β-cardiac myosin to form the IHM, confirming now a previous suggestion based on the hypothesis that the heads would remain in the PPS state while forming the IHM^46^.

## Mutations in the lever arm helix and the light chains

Studies of the mutations in the light-chains and the lever arm have been performed (reviewed by ^43, 45^). HCM causing mutations in the light chains are generally considered as rare: 1% in the RLC and <1% in the ELC^57^. Mutations in the heavy chain from the Pliant and IQ regions were also recently studied: **D778V**, **L781P** and **S782N** in the Pliant; **A797T** in IQ1 and **F834L** in IQ2^58^ (**Extended data Fig. 10b, 10c**). Importantly, none of these mutations have effects on light-chain loading to the heavy chain. The three Pliant mutations have no significant effect for the SRX but alter the mechano-chemical parameters of the myosin^58^. In contrast, mutations **A797T** and **F834L** have modest or no effect on mechano-chemical parameters but significantly reduce the SRX^58^. The replacement of the small **Ala797** by a bulkier Threonine is predicted to add restrains on the orientation of _ELC_α2-helix and may influence the orientation dynamics of the entire N-terminal lobe of the ELC, thus weakening the ELC/RLC interface by long-range effects. In IQ2, the replacement of the aromatic **Phe834** with the smaller Leu could have effects on the RLC/RLC interface and may alter the local orientation of the helices involved in this interface, thus weakening it. Both of these mutations in the IQs are predicted to have long-range effects as they would impact the formation of the hinges required in the lever arm for the formation of the IHM.

## Mutations in the S2 coiled-coil

Finally, the effect of HCM mutations located in the S2 coiled-coil on the SRX has not yet been investigated. However, the effect of two mutations on the affinity of the S1 for the S2 was evaluated: **R870H** and **D906G**^4^. While **R870H** has no effect on the association of the S1 and S2, D906G reduces the affinity^4^. In our IHM structure, the R870H is located far from the interface and would not affect the coiled-coiled helices association (**Extended data Fig. 9a**). Modeling of the coiled coil indicates that this is not the case for the D906G mutation which could be relatively close to the interface. The addition of a Gly could weaken the helical secondary structure of the S2 coiled-coil and thus weaken the IHM interfaces with long-range repercussions. A recent study^59^ indicates in fact that R870H has an increase in affinity for phosphorylated MyBP-C, which could lead to the HCM disease by preventing the proper role of phosphorylated MyBPC-C as well as destabilizing heads in the IHM when MyBP-C is phosphorylated.

## Conclusions

The structure of the IHM at high-resolution solved in this work enables prediction and analysis of the effect of point mutations that cause HCM. It is an essential tool to understand the molecular basis of cardiomyopathy development. As proposed previously, the point mutations could have different effects on motor function and/or stability of the sequestered state ^46^. Interestingly, the mutations altering the stability of the sequestered state can be classified in different categories: **(i)** those directly altering the interface; **(ii)** mutations with indirect effect on the interface, destabilizing elements involved in the interfaces; **(iii**) mutations located in connectors and subdomains essential to stabilize states with lever arm up, with long range effects. The structure will be of high utility to discuss the outcome of HCM and DCM mutations. Structural studies and dynamic simulations of mutants can be performed based on the protocol or structure this study has provided. This will be essential to reveal how mutations affect force production and the stability of the auto-inhibited state. This knowledge will be beneficial to the patient, as predicting the effect mutations have on motor activity or SRX stability will be critically needed as different types of allosteric modulators of cardiac myosin function become available.

**Extended data Movie 1:** overview on the map 1 (EMD-15353) at a global resolution of 3.6 Å. The blocked head is colored in blue, the free head is colored in green. The ELCs are colored in pink and the RLCs are colored in light brown. Different views are represented, the final view displays the head-head interface with _FH_Top-Loop (orange), _BH_Loop-4 (cyan) and _BH_HCM-loop (red).

**Extended data Movie 2:** overview on the map 2 (EMD-15354) at a global resolution of 3.2 Å. The blocked head is colored in blue, the free head is colored in green. The ELCs are colored in pink and the RLCs are colored in light brown. Different views are represented. In the views of the active site, Switch-1 is colored in black, Switch-2 is colored in orange, P-loop is colored in purple. The final view displays the head-head interface with _FH_Top-Loop (orange), _BH_Loop-4 (cyan) and _BH_HCM-loop (red).

**Extended data Movie 3:** Illustration of the asymmetry of the RLC-RLC interface. See **Extended data Fig. 7**.

---

**Extended data Figure 1.**
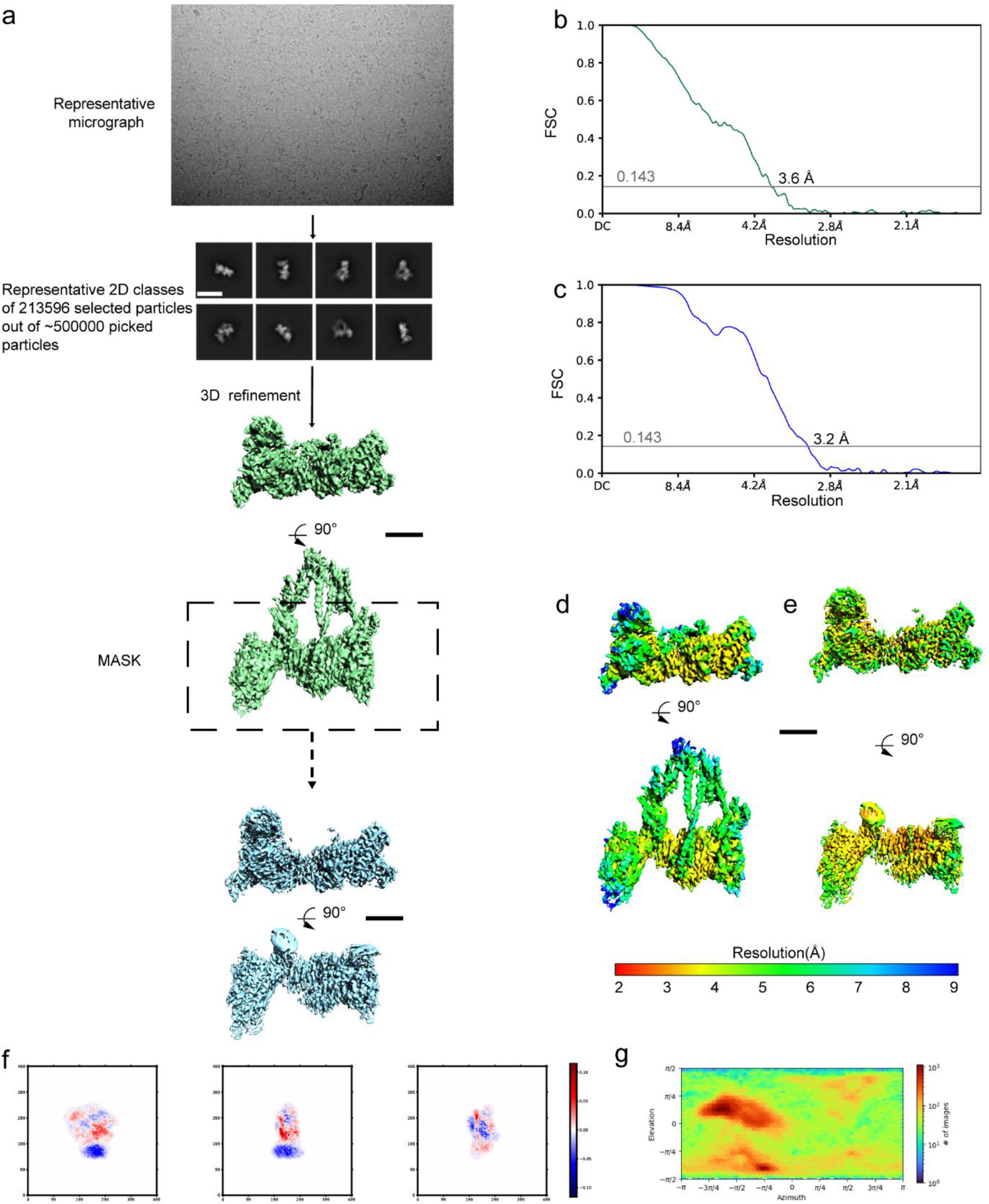
– CryoEM map and validation. **(a)** Shows the steps of processing. A representative micrograph on gold grids, 2D classes and the steps of refinement and masking. **(b)** and **(c)** show the Fourier Shell Correlation and the overall resolution on Map 1 and 2 respectively (criterion 0.143). **(d)** and **(e)** display the local resolution for each map with two orientations. **(e)** Shows the flexibility within the different classes used to reconstitute the map. **(f)** plots the different orientations of the particles.

**Extended data Figure 2.**
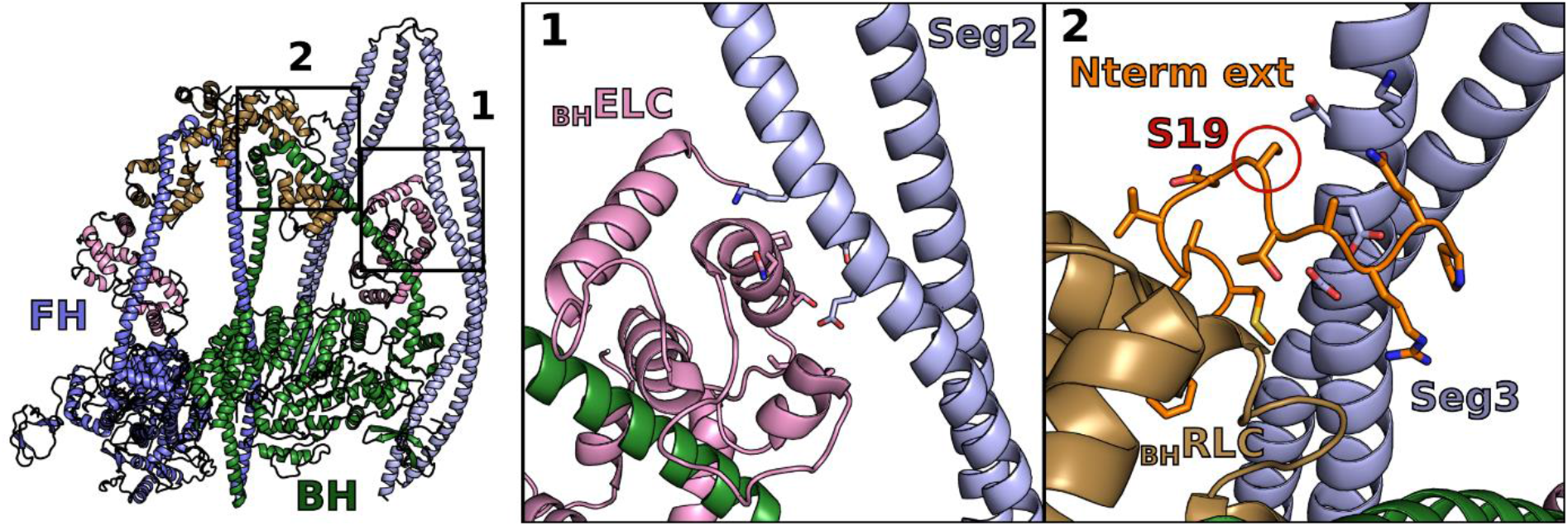
– Segments 2 (Seg2) and 3 (Seg3) coiled-coils provide additional stability in the Smooth muscle myosin 2 (SmMyo2) IHM structure. Seg2 interacts with the ELC from the blocked head (BH) (interface 1) and Seg3 with the N-terminal extension (N-term ext) from the RLC from the BH (interface 2). The phosphorylatable serine (S19) in the _BH_RLC is part of the interface with the Seg3. IHM model used here: PDB code 7MF3^1^.

**Extended data Figure 3.**
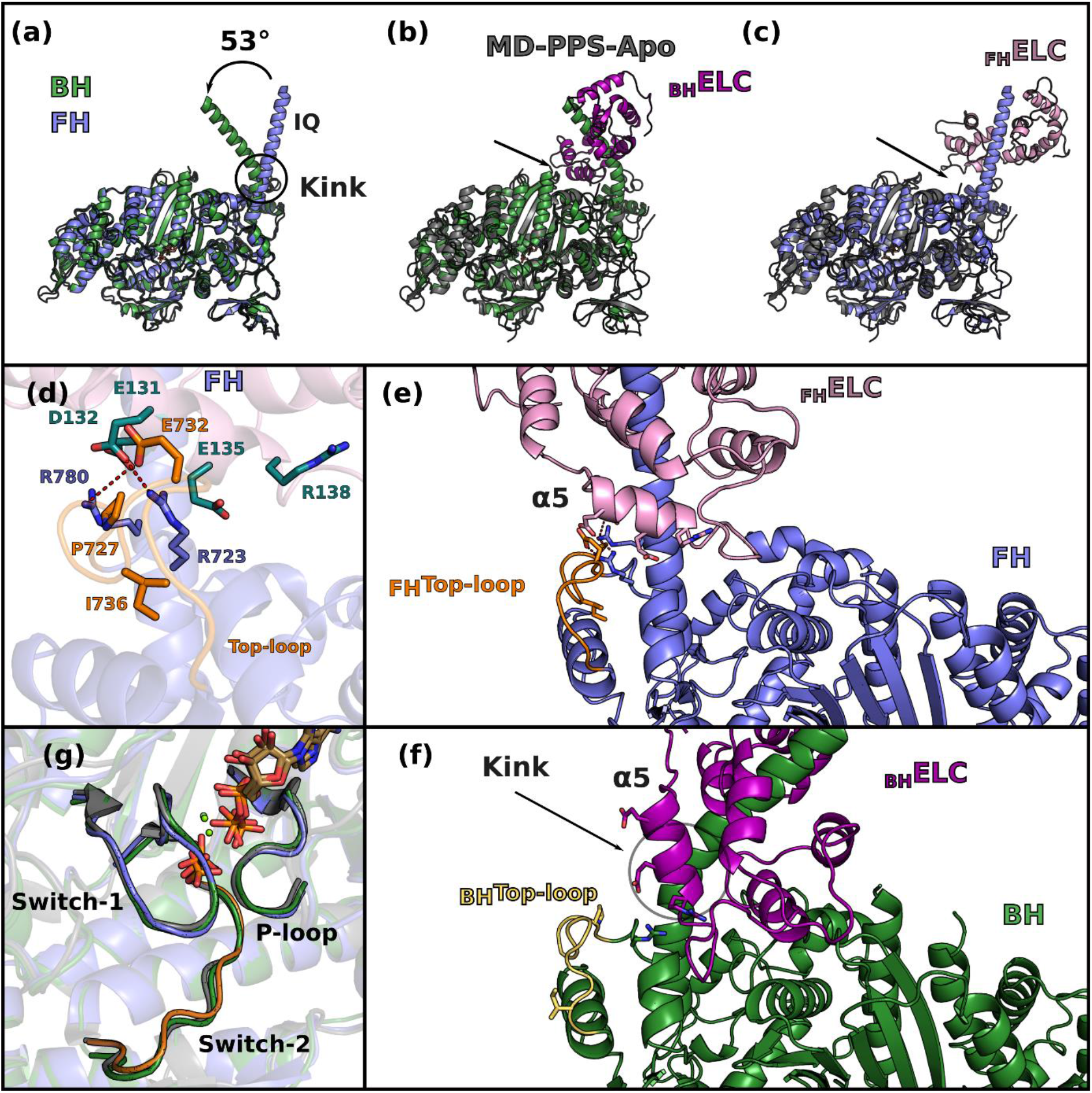
– Comparison of the sequestered state with the pre-powerstroke (PPS) state of cardiac myosin. **(a)** The blocked head (BH) and the free head (FH) superimpose on the motor domain with a RMSD of 0.7 Å, the only difference being the kink at the Pliant region in the lever arm, resulting in an angle of 53° between the IQ1 helix of the BH and the FH heads. **(b)** and **(c)** show the superimposition of the BH and of the FH respectively with the apo cardiac MD structure in the PPS state (black lines, MD-PPS-Apo, PDB code 5N6A,^2^). Both motor domains superimpose with MD-PPS-Apo with a rmsd of 0.84 Å. The arrows show the ELC/Converter interface which is altered by the kink in the BH. **(d)** In the FH, the Converter/ELC interface is maintained by “musical chairs”, a set of labile charged residues located on the ELC (deep teal cyan) or in the Converter, more specifically in the Top-loop^3^ (orange). **(e)** and **(f)** compare the motor/ELC interface and how it is completely altered by the kink present in the BH, the musical chairs are represented on both panels. The kink of the BH shifts the position of the helix. In the BH, the interface is altered by the kink, generating a new interface between the motor (green with Top-Loop in light orange) and the ELC (purple). **(g)** BH, FH and PPS-S1-OM have all a closed active site in the inner cleft region.

**Extended data Figure 4.**
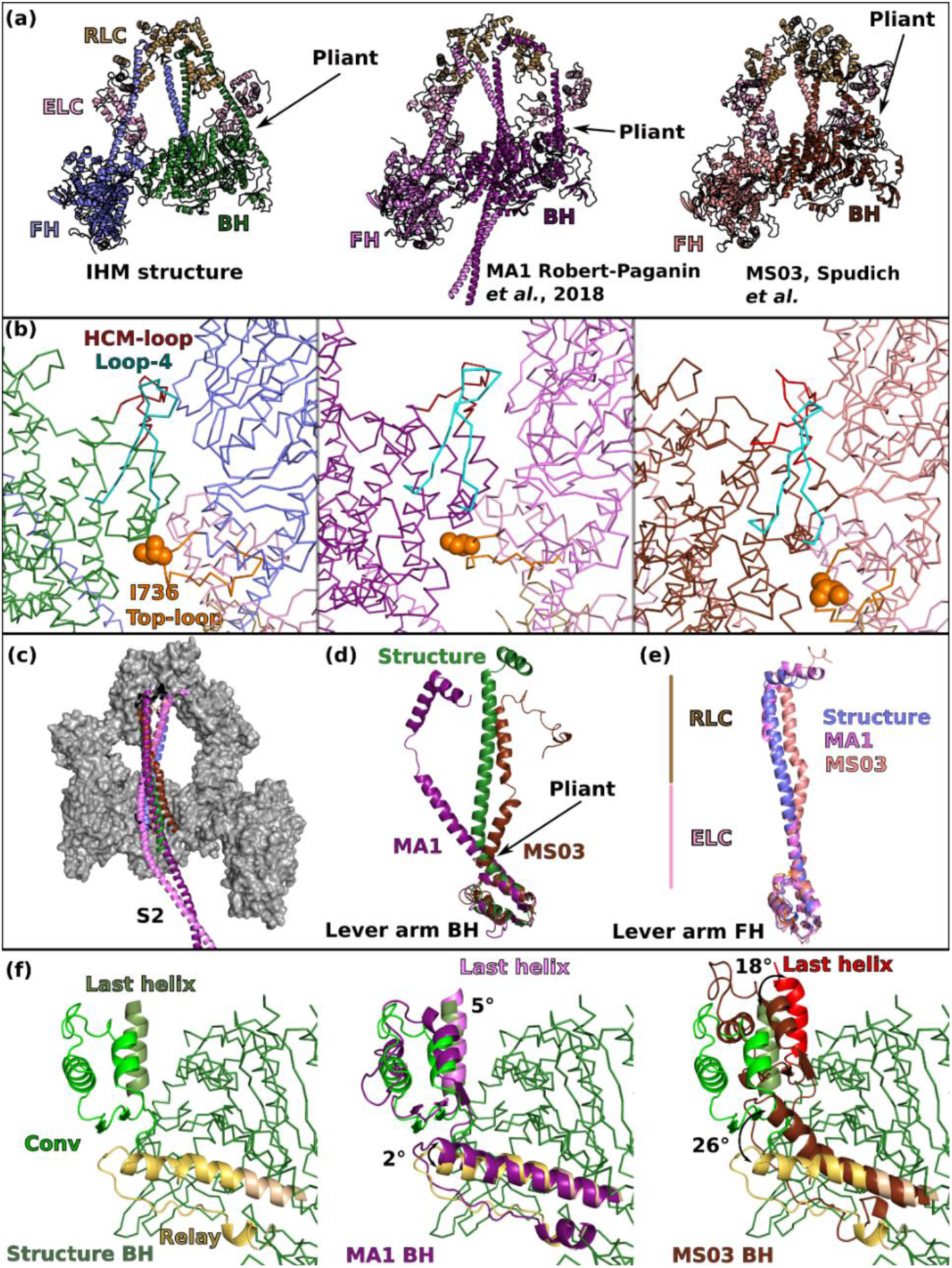
– Comparison of the IHM structure with the previous homology pseudo-atomic models. **(a)** Overall representation of the IHM structure of cardiac myosin with two models, MA1^3^ and MS03^4^. The pliant region of the BH is shown with an arrow. **(b)** Zoom in the head-head interface region showing that the same elements are involved in the interface, although the interface was not correctly modeled in the details. **(c)** S2 coiled-coil in the structure and in the models interacts with the BH. **(d)** and **(e)** compare the conformation of the lever arm of the BH and the FH respectively in the structure and in the models. The regions where the ELC and the RLC bind are indicated. **(f)** Comparison of the orientation of the Relay and the Converter in the BH. The structure (left) is superimposed with the two models (middle, right) for comparison. Note the pre-prestroke position of the Relay (brown) and the Converter (brown, red) in the MS03 model. In **(a)** and **(b)**, the structures are superimposed on the entire model, in (**c**), the structures are superimposed on both heads (which include both Motor domains and the lever arms including the two LCs), in **(d)** and **(e)** the lever arms are superimposed on the Converter (residues 708-777), in **(f)**, the structures are superimposed on the N-term subdomain (residues 3-202).

**Extended data Figure 5.**
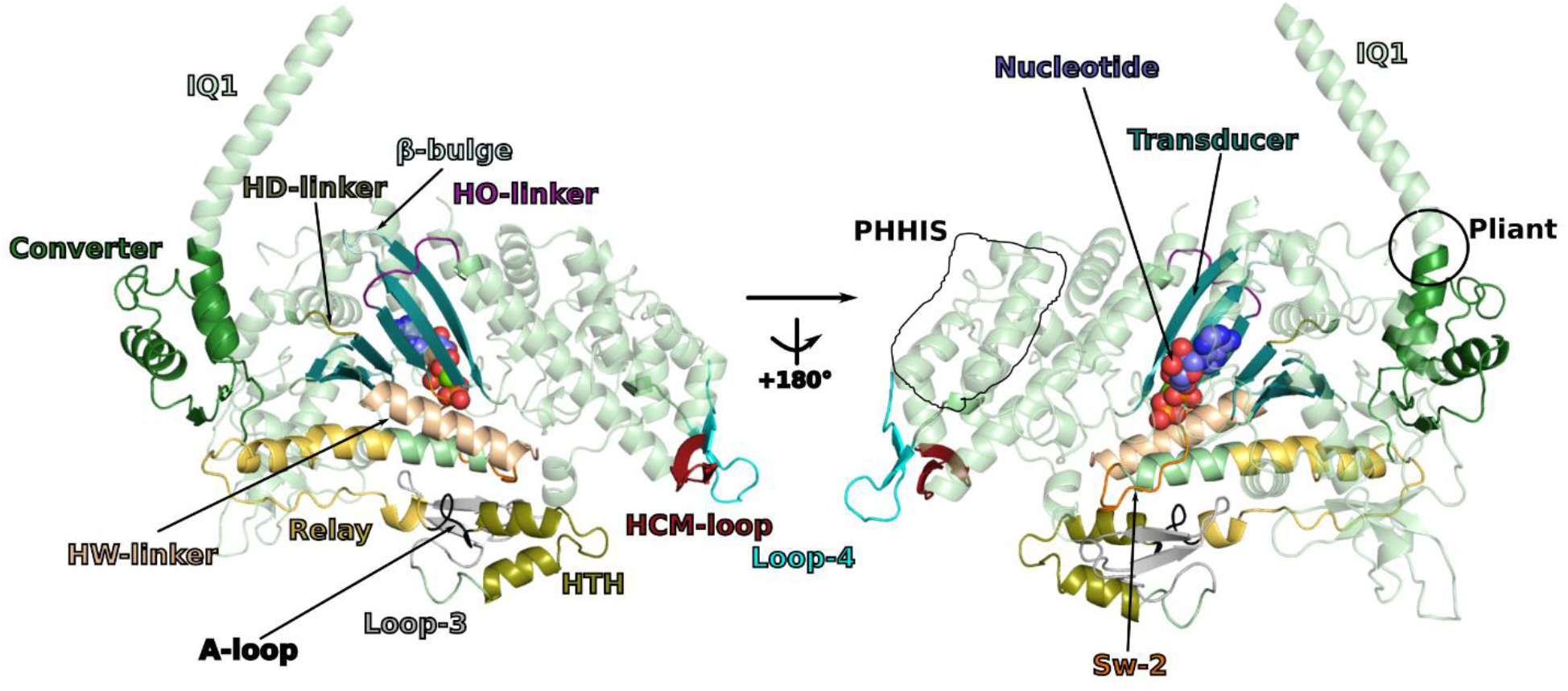
– Elements and connectors of β-cardiac myosin involved in the interfaces stabilizing the IHM. The different elements are represented on the blocked head of cardiac IHM. The nucleotide indicates the position of the active site while elements of the actin interface are represented : HTH: helix-turn-helix, A-loop: activation-loop, HCM-loop, Loop-4 and Loop-3. Also represented are Sw-2 (switch-2), IQ1 helix: HC sequence involved in binding the ELC after the pliant region. The Transducer (blue) corresponds to the central β-sheet as well as structural elements linked to this β-sheet (β-bulge, HO-linker, HD-linker). The PHHIS is part of the U50 subdomain while the HTH and Loop-3 are part of the L50 subdomains.

**Extended data Figure 6.**
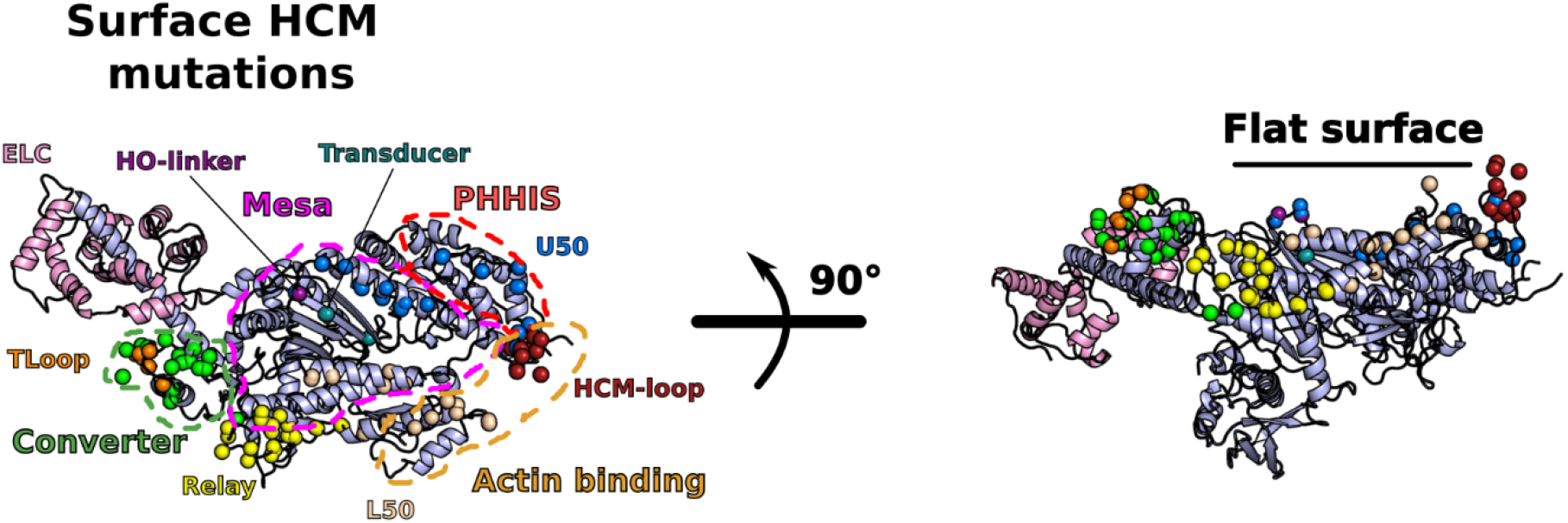
– Definition of the myosin “Mesa” and elements involved in the interfaces of the IHM, as defined by^5^. Surface HCM mutations are shown as balls and colored depending on the subdomain where they are located. The Mesa was defined as a large and flat surface containing conserved residues and HCM mutations (in purple dotted lines). The Primary Head-Head Interaction Site (PHHIS) and the actin-binding surface are shown with red and orange dotted lines respectively.

**Extended data Figure 7.**
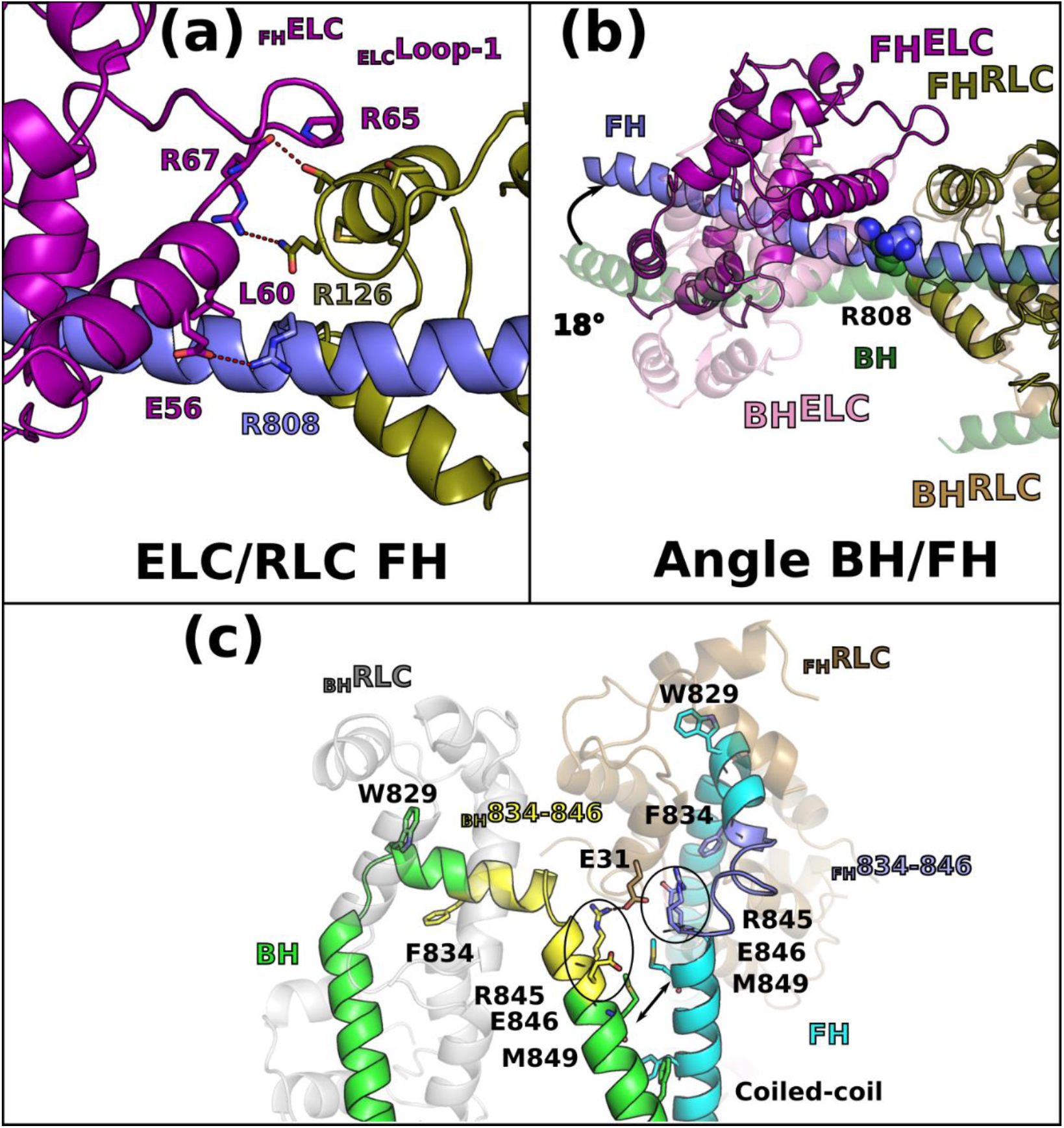
– Asymmetry in the light chain binding regions. **(a)** Interactions at the ELC/RLC interface in the free head (FH) differ from the ELC/RLC interface in the blocked head (BH) (Fig. 2d). **(b)** Lever arm of the BH and of the FH aligned on IQ2 reveals a 18° difference in the orientation as well as major differences in the ELC/RLC interface**. (c)** Asymmetry in the conformation and contacts of the two RLCs in the IHM structure. The RLC/IQ2 of the BH and FH heads see a different environment. This is specifically true for the region 834-848 of the heavy chain (HC), due to the fact that the coiled-coil triggers a shift in position of the HC residues (arrows between the M849 position). The HC residues (834-848) thus adopt drastically different conformations (yellow for BH, dark blue for FH) that are critical to form the RLC/RLC interface. This shift explains why some residues have different environments in the FH and in the BH: two residues characteristic of this phenomenon are contoured in black, R845 and E846. After residue R846, the phase of the coiled-coil is more canonical. Interestingly, the _BH_R845 residue is involved in electrostatic interactions with the _FH-RLC_E31 residue, while the _FH_R845 residue is buried in the _FH-RLC_N-lobe core. See also ***Movie 3***.

**Extended data Figure 8.**
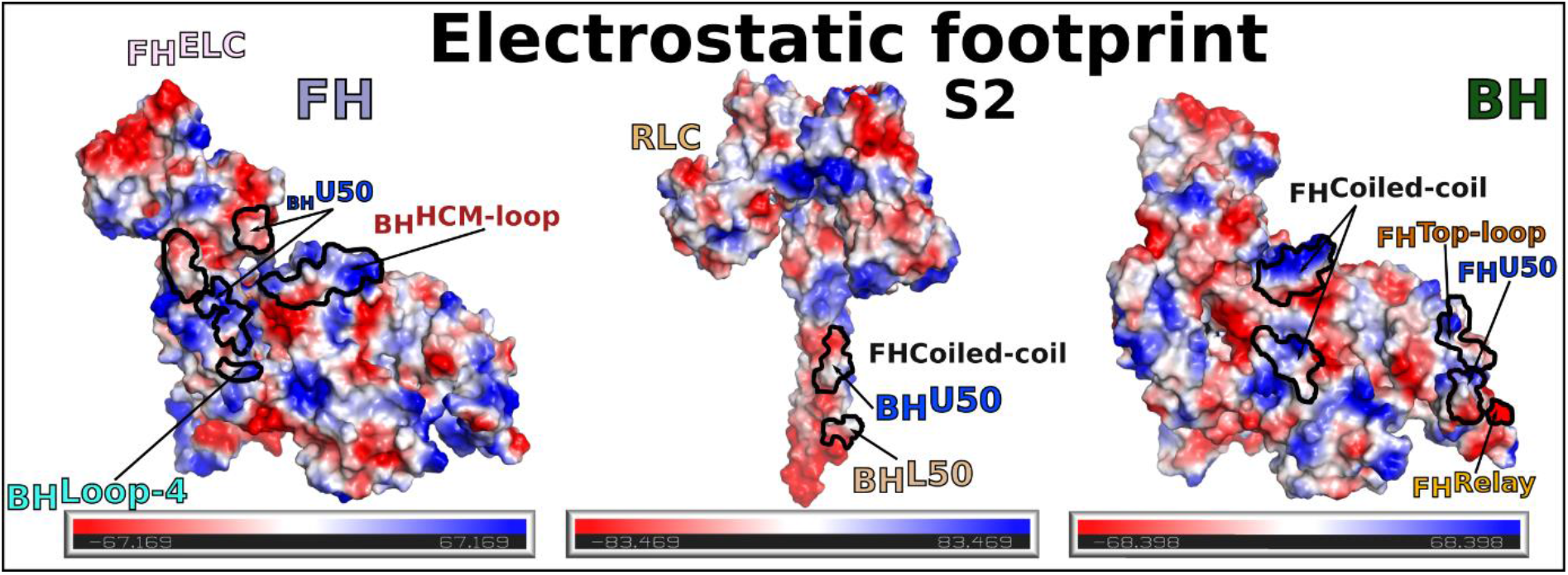
– Electrostatic profile of the footprint of IHM interfaces. The different surfaces involved in the interactions stabilizing the IHM are represented as electrostatic surfaces. The regions involved in the interactions are indicated.

**Extended data Figure 9.**
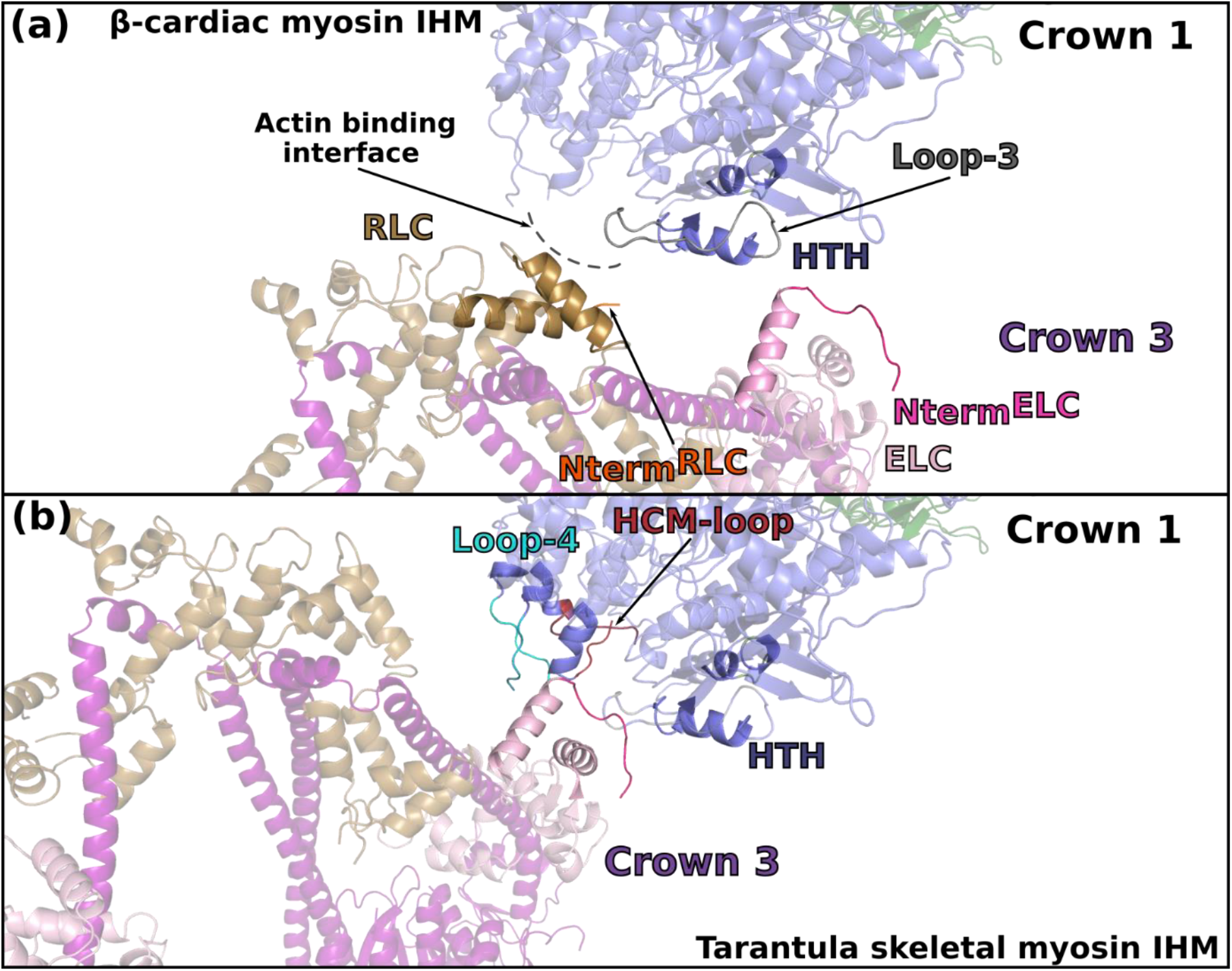
– Comparison of the inter-crown interfaces between human β-cardiac myosin IHM. **(a)** and tarantula skeletal myosin IHM (**b**). In order to make the comparison, two IHM of β-cardiac myosin were fitted in the low-resolution map of tarantula skeletal myosin IHM (EMDB-1950^6^). The difference in the symmetry of cardiac and tarantula skeletal filaments (pseudo-helical and purely helical respectively) induces different inter-crown interfaces. In tarantula skeletal relaxed filament, the ELC from Crown 3 interacts with elements of the actin binding interface from the adjacent Crown 1: Loop-4, HCM-loop and helix-turn-helix (HTH). There is no contribution of the RLC in the inter-crown interface. In β-cardiac myosin, the inter-crown interface involves the both the ELC and the RLC from Crown 3 and HTH and Loop-3 from Crown 1. These differences in the interfaces are likely to be responsible of the distinct regulatory mechanisms of the two types of filaments.

**Extended data Figure 10.**
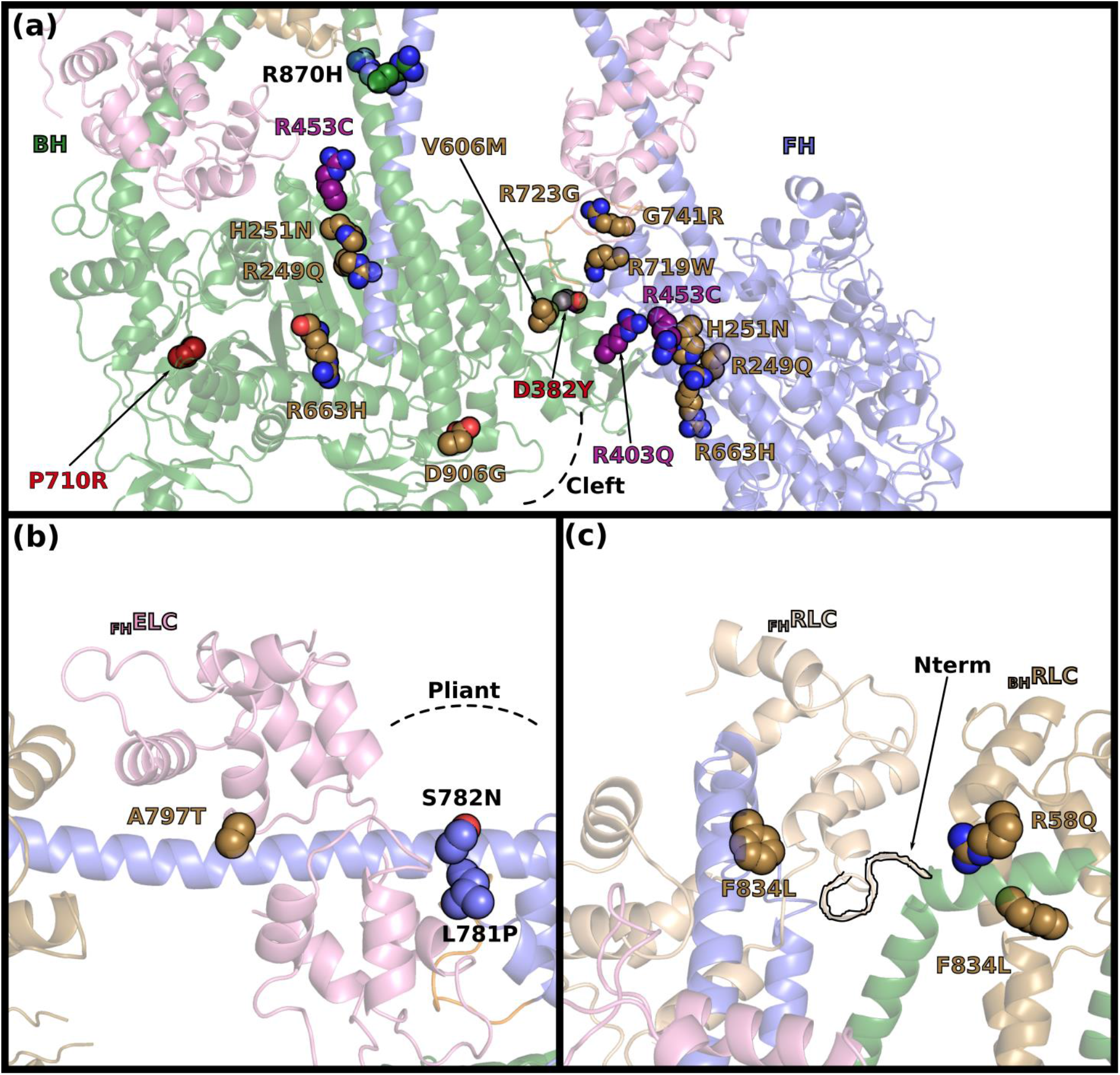
– Effects of HCM mutations on myosin function and regulation. A set of HCM mutations with effect on the SRX or stability of the interfaces are shown with side chains as spheres. Mutations with direct destabilization of the interface are colored in purple; mutations with indirect effect on the interface are colored in yellow sand; mutation without effect on the stability of the IHM are not colored (name in black); mutations with long range effects and major effects on myosin structural changes are colored in red. **(a)** Head/head and coiled-coiled/head interfaces. P710R is located at the base of the Converter. The internal cleft close to the actin-binding interface (cleft) is shown with dashed lines. **(b)** Mutations in the Pliant region and in the ELC. **(c)** Mutations at the RLC/RLC interface, the N-terminal extension (Nterm) has been contoured in black.

**Extended data Table 1:** EM validation

|  |  |
| --- | --- |
| **IHM β-cardiac myosin (EMD-15353)** |  |
| **Data collection and processing** |  |
| Microscope | Titan KRIOS G3 |
| Detector | K3 |
| Magnification | 105k |
| Total exposure (e-/Å<sup>2</sup>) | 39.2 |
| Exposure time | 2.4 s |
| Electron exposure per frame (e-/Å<sup>2</sup>) | 0.98 |
| Defocus range (μm) | -0.8, -2.2 |
| Pixel size (Å) | 0.84 |
| Symmetry imposed | C1 |
| Initial particles images (no.) | 493179 |
| Final particle images (no.) | 213596 |
| Map resolution (Å) | 3.6 |
| FSC threshold | 0.143 |
| Map resolution range (overall) | 3-9 |
| **Refinement** |  |
| Initial model used (PDB codes) | 6N69, 2XFM, homology modeling from 7MF3 |
| Map sharpening B-factor (Å<sup>2</sup>) |  |
| **Model composition** |  |
| Non-H atoms | 18839 |
| Protein residues | 2322 |
| Ligands | 6 |
| *RMSD from ideal geometry* |  |
| Bond length (Å) | 0.002 |
| Bond angles (°) | 0.601 |
| **Validation** |  |
| Molprobit score | 1.87 |
| clashcore | 16.05 |
| *Ramachandran (%)* |  |
| Outliers | 0 |
| Allowed | 2.89 |
| Favored | 97.11 |
| *Validation* |  |
| CC (mask) | 0.79 |
| CC (volume) | 0.78 |
| EMRinger score (overall mode) | 0.81 |
| EMRinger score (1-786, heads regions at high resolution) | 1.15 |
| **IHM β-cardiac myosin head-head masked map (EMD-15354)** |  |
| **Data collection and processing** |  |
| Microscope | Titan KRIOS G3 |
| Detector | K3 |
| Magnification | 105k |
| Total exposure (e-/Å<sup>2</sup>) | 39.2 |
| Exposure time | 2.4 s |
| Electron exposure per frame (e-/Å<sup>2</sup>) | 0.98 |
| Defocus range (μm) | -0.8, -2.2 |
| Pixel size (Å) | 0.84 |
| Symmetry imposed | C1 |
| Initial particles images (no.) | 493179 |
| Final particle images (no.) | 213596 |
| Map resolution (Å) | 3.2 |
| FSC threshold | 0.143 |
| Map resolution range (overall) | 2.5-6 |
| CC (mask) | 0.61 |
| CC (volume) | 0.61 |
| EMRinger score (1-786, heads regions at high resolution) | 1.78 |

**Extended data Table 2:**
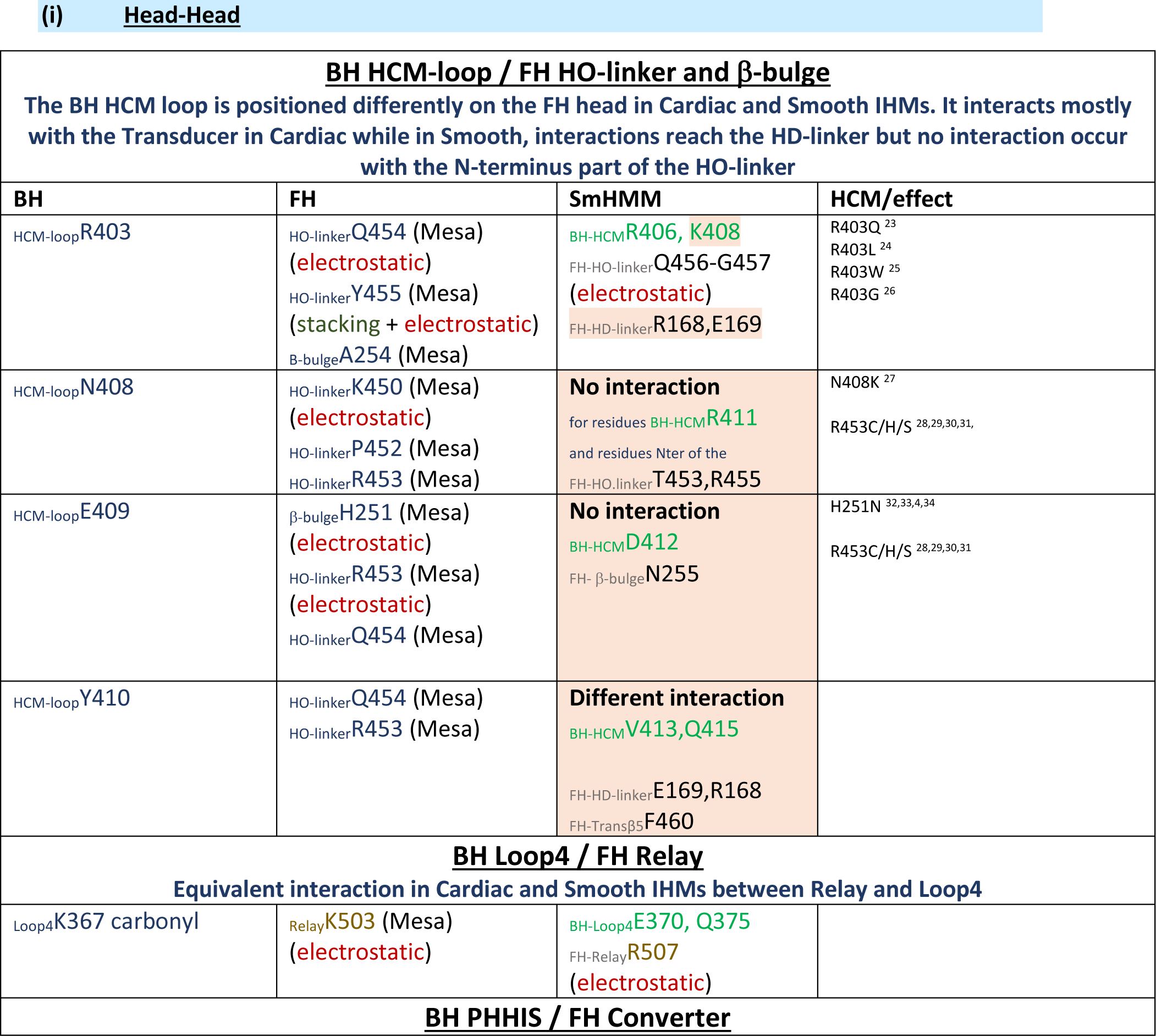
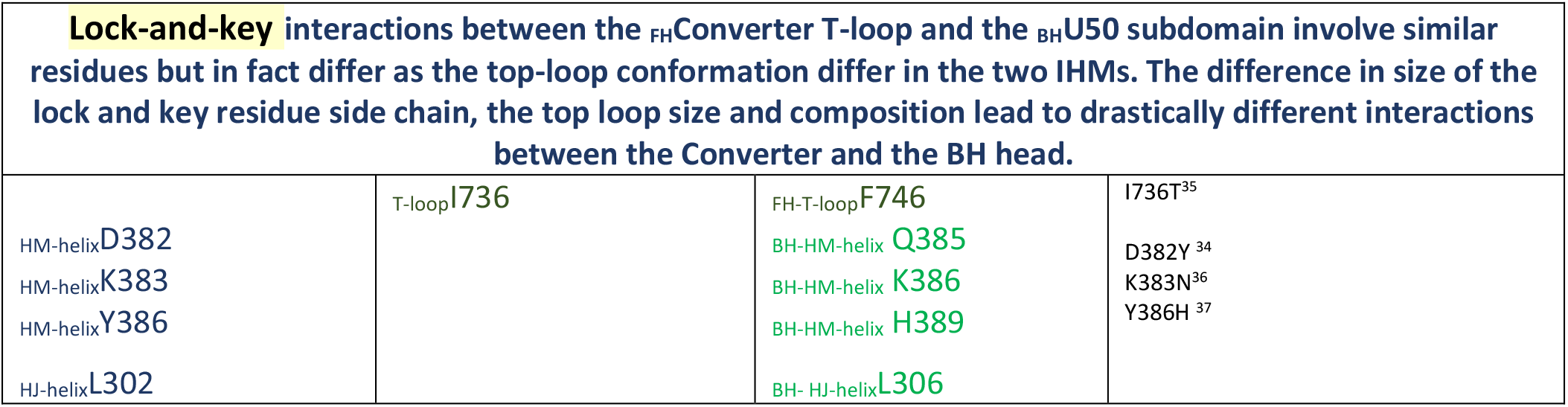
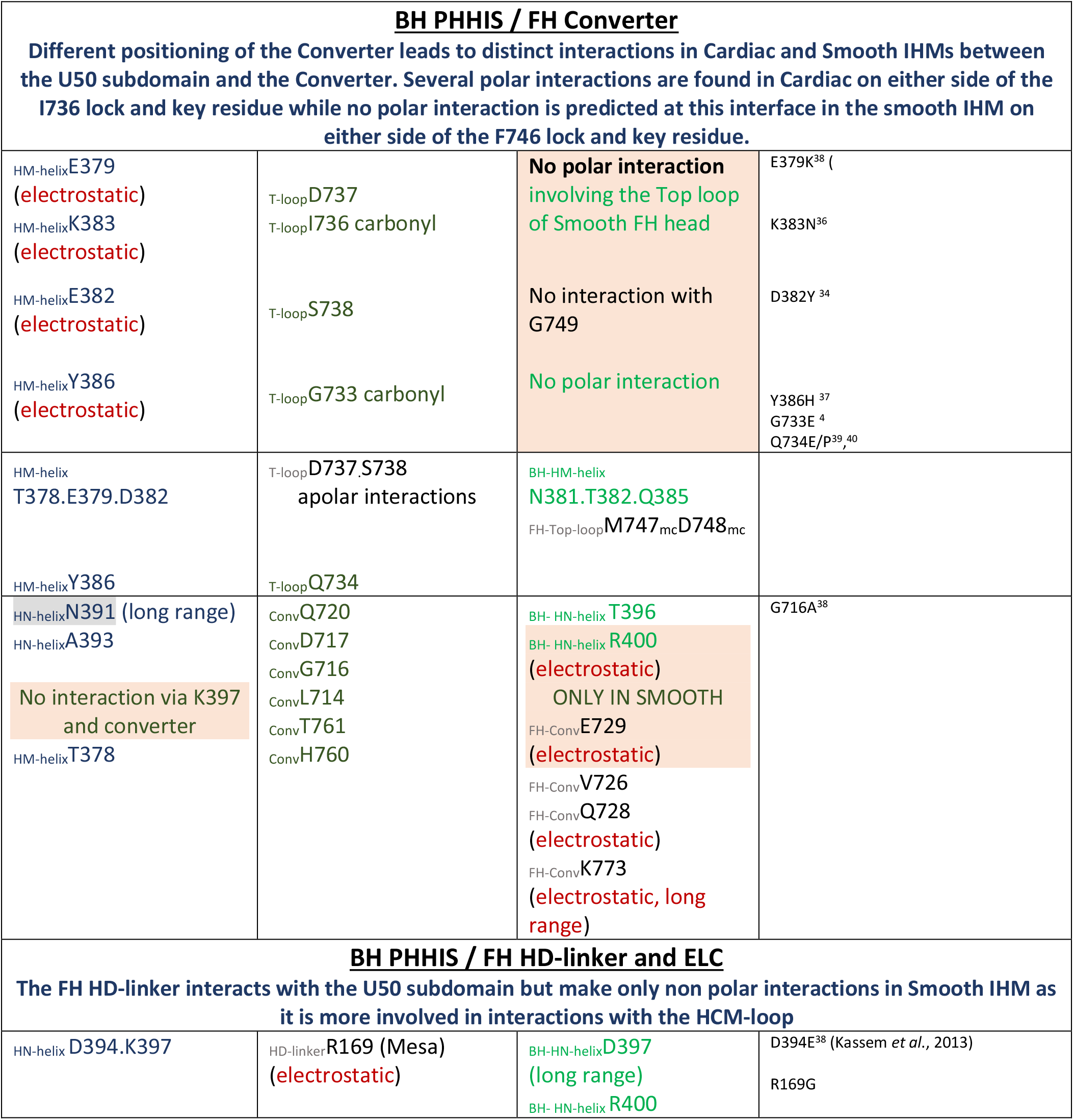
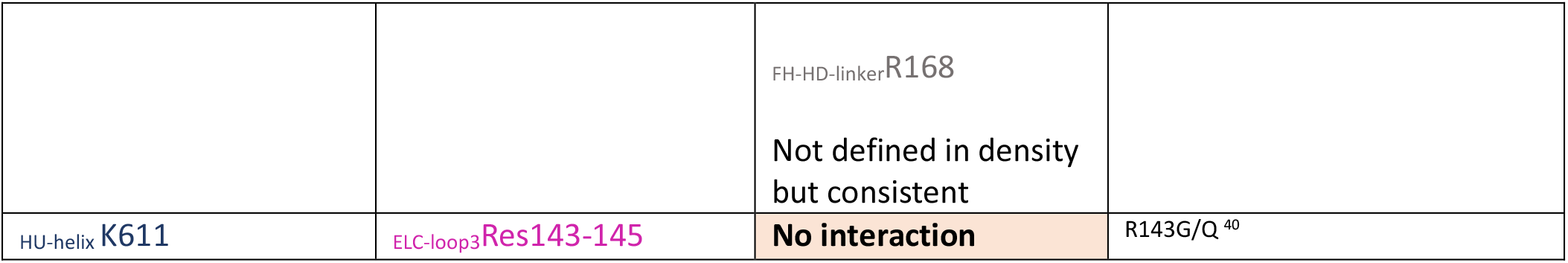
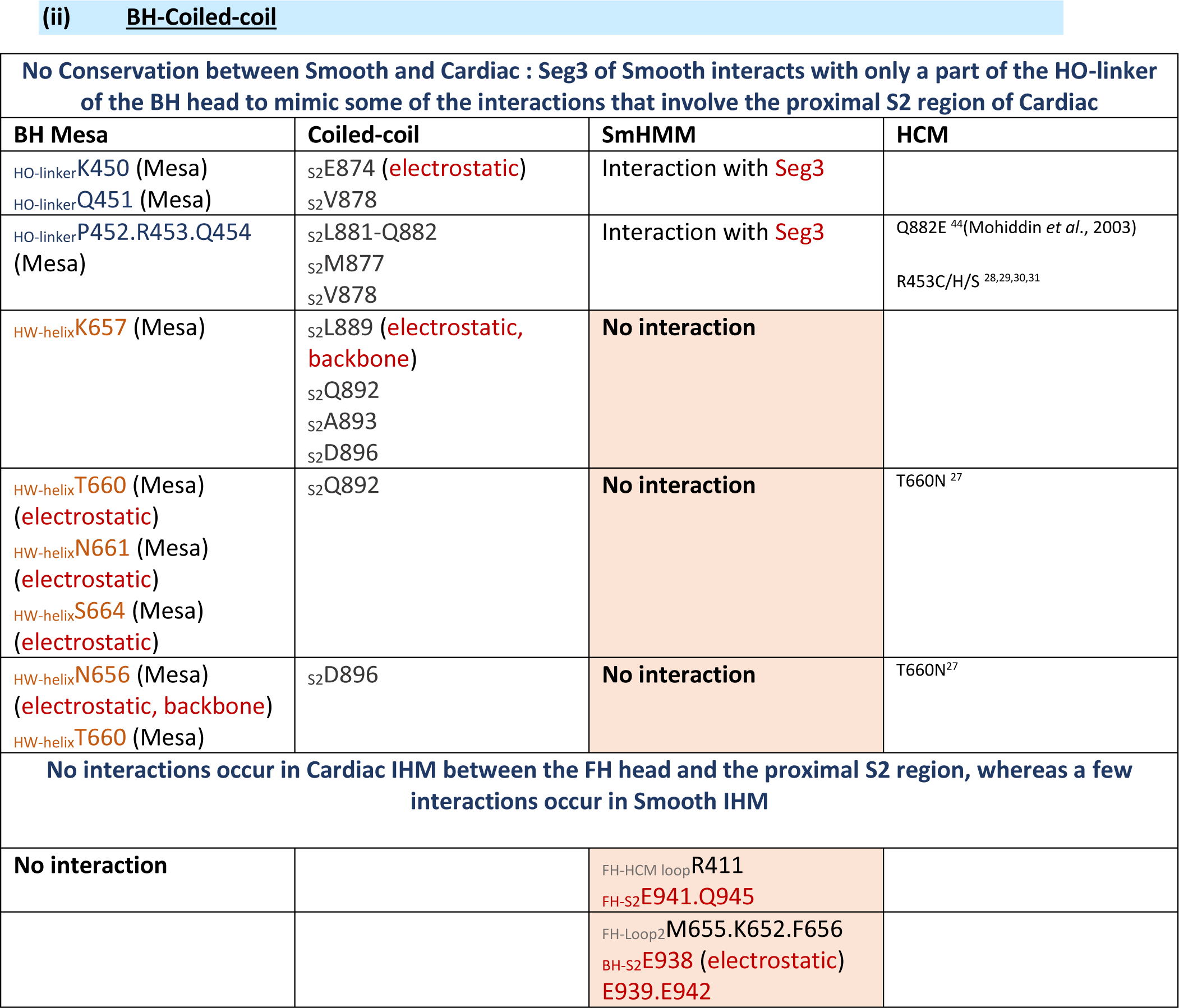
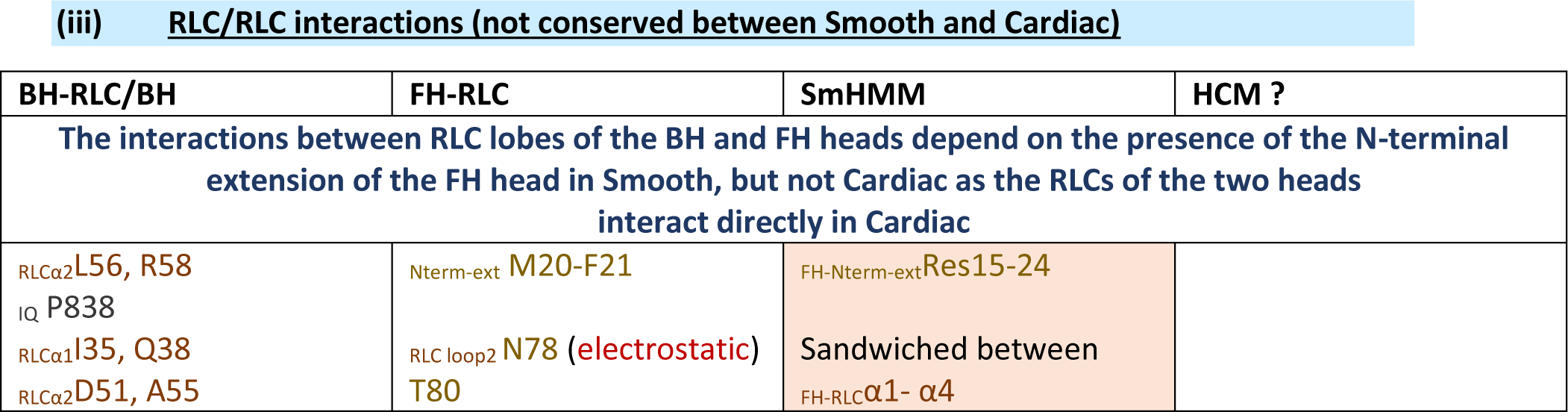
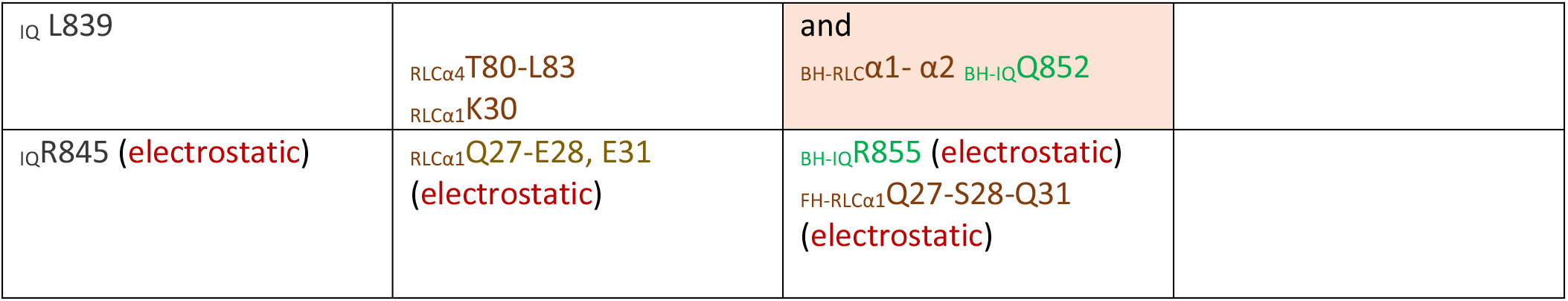
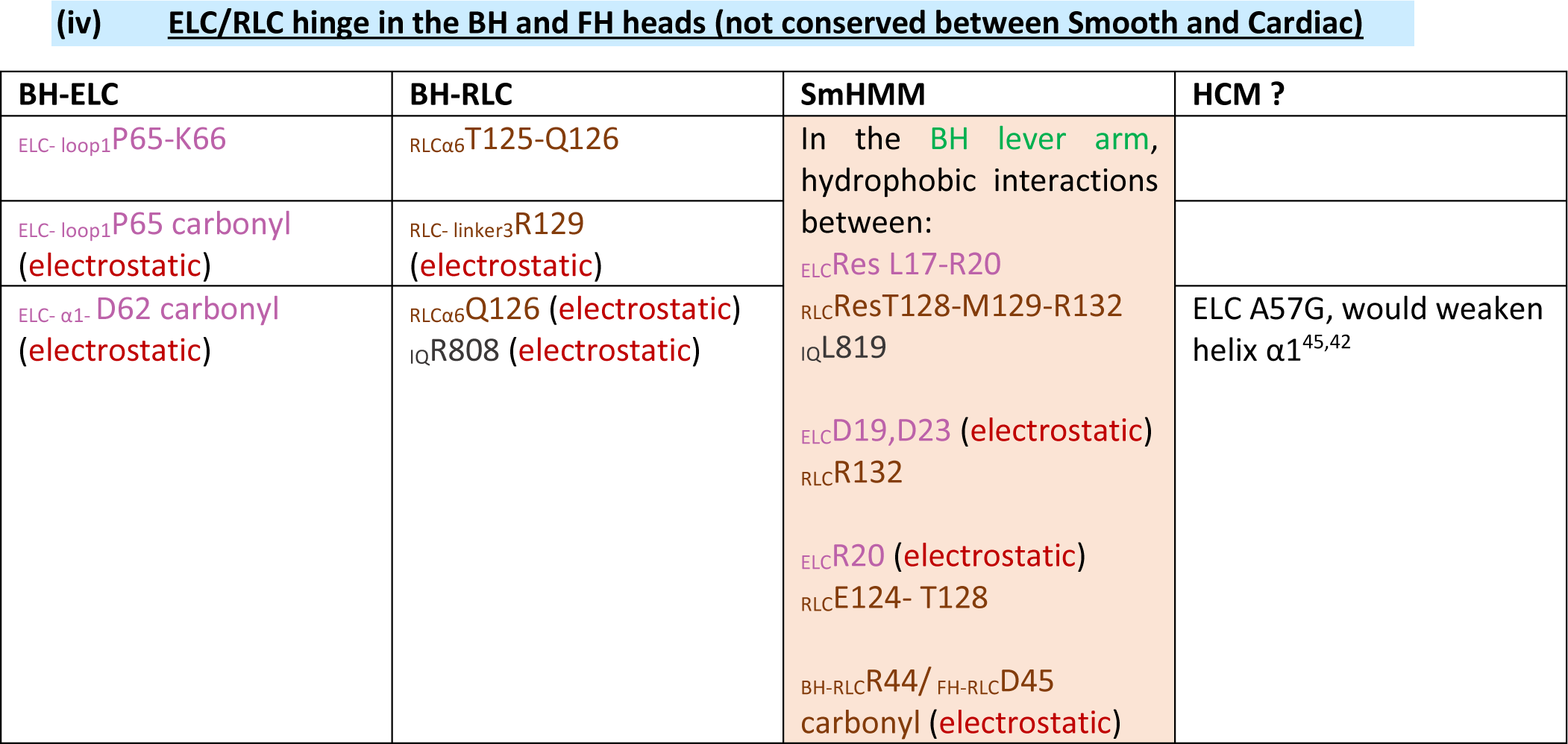
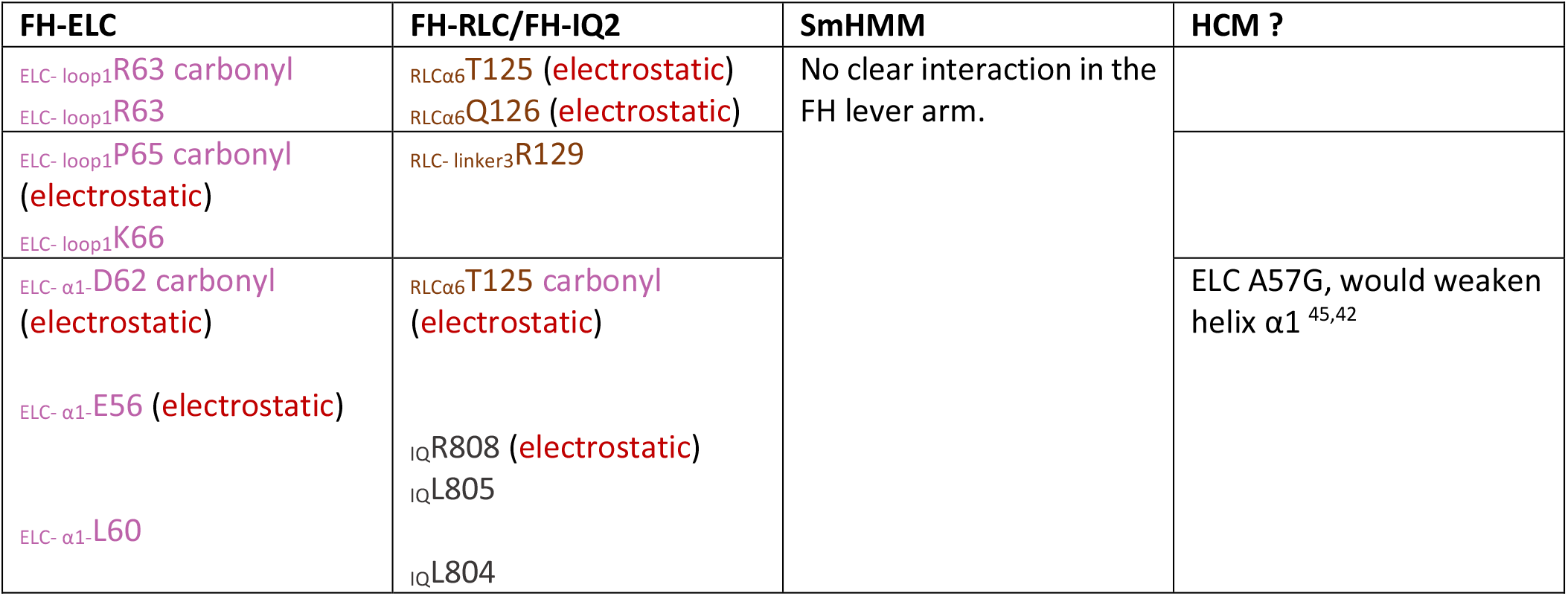
Interactions stabilizing cardiac IHM: Interactions that differ drastically between smooth and cardiac are shown with orange background.

## References

1. Robert-Paganin, J., Pylypenko, O., Kikuti, C., Sweeney, H. L. & Houdusse, A. Force Generation by Myosin Motors: A Structural Perspective. Chem. Rev. 120, 5–35 (2020).

2. Janssen, I., Heymsfield, S. B., Wang, Z. M. & Ross, R. Skeletal muscle mass and distribution in 468 men and women aged 18-88 yr. J. Appl. Physiol. 89, 81–88 (2000).

3. Stewart, M. A., Franks-Skiba, K., Chen, S. & Cooke, R. Myosin ATP turnover rate is a mechanism involved in thermogenesis in resting skeletal muscle fibers. Proc. Natl. Acad. Sci. 107, 430–435 (2010).

4. Naber, N., Cooke, R. & Pate, E. Slow Myosin ATP Turnover in the Super-Relaxed State in Tarantula Muscle. J. Mol. Biol. 411, 943–950 (2011).

5. Hooijman, P., Stewart, M. A. & Cooke, R. A new state of cardiac myosin with very slow ATP turnover: A potential cardioprotective mechanism in the heart. Biophys. J. 100, 1969–1976 (2011).

6. Schmid, M. & Toepfer, C. N. Cardiac myosin super relaxation (SRX): a perspective on fundamental biology, human disease and therapeutics. Biol. Open 10, (2021).

7. Cross, R. A., Jackson, A. P., Citi, S., Kendrick-Jones, J. & Bagshaw, C. R. Active site trapping of nucleotide by smooth and non-muscle myosins. J. Mol. Biol. 203, 173–181 (1988).

8. Wendt, T., Taylor, D., Trybus, K. M. & Taylor, K. Three-dimensional image reconstruction of dephosphorylated smooth muscle heavy meromyosin reveals asymmetry in the interaction between myosin heads and placement of subfragment 2. Proc. Natl. Acad. Sci. 98, 4361–4366 (2001).

9. Woodhead, J. L. et al. Atomic model of a myosin filament in the relaxed state. Nature 436, 1195–1199 (2005).

10. Al-Khayat, H. A., Kensler, R. W., Squire, J. M., Marston, S. B. & Morris, E. P. Atomic model of the human cardiac muscle myosin filament. Proc. Natl. Acad. Sci. U. S. A. 110, 318–23 (2013).

11. Lee, K. H. et al. Interacting-heads motif has been conserved as a mechanism of myosin II inhibition since before the origin of animals. Proc. Natl. Acad. Sci. U. S. A. 115, E1991–E2000 (2018).

12. Alamo, L., Pinto, A., Sulbarán, G., Mavárez, J. & Padrón, R. Lessons from a tarantula: new insights into myosin interacting-heads motif evolution and its implications on disease. Biophys. Rev. 10, 1465–1477 (2018).

13. Scarff, C. A. et al. Structure of the shutdown state of myosin-2. Nature 588, 515–520 (2020).

14. Yang, S. et al. Cryo-EM structure of the inhibited (10S) form of myosin II. Nature 588, 521–525 (2020).

15. Heissler, S. M., Arora, A. S., Billington, N., Sellers, J. R. & Chinthalapudi, K. Cryo-EM structure of the autoinhibited state of myosin-2. Sci. Adv. 7, eabk3273 (2021).

16. Brunello, E. et al. Myosin filament-based regulation of the dynamics of contraction in heart muscle. Proc. Natl. Acad. Sci. U. S. A. 117, 8177–8186 (2020).

17. Adhikari, A. S. et al. β-Cardiac myosin hypertrophic cardiomyopathy mutations release sequestered heads and increase enzymatic activity. Nat. Commun. 10, 2685 (2019).

18. Spudich, J. A. The myosin mesa and a possible unifying hypothesis for the molecular basis of human hypertrophic cardiomyopathy. Biochem. Soc. Trans. 43, 64–72 (2015).

19. Spudich, J. A. et al. Effects of hypertrophic and dilated cardiomyopathy mutations on power output by human beta-cardiac myosin. J Exp Biol 219, 161–167 (2016).

20. Spudich, J. A. Three perspectives on the molecular basis of hypercontractility caused by hypertrophic cardiomyopathy mutations. Pflugers Arch. 471, 701–717 (2019).

21. Robert-Paganin, J., Auguin, D. & Houdusse, A. Hypertrophic cardiomyopathy disease results from disparate impairments of cardiac myosin function and auto-inhibition. Nat. Commun. 9, 4019 (2018).

22. Kampourakis, T., Zhang, X., Sun, Y.-B. & Irving, M. Omecamtiv mercabil and blebbistatin modulate cardiac contractility by perturbing the regulatory state of the myosin filament. J. Physiol. 596, 31–46 (2018).

23. Anderson, R. L. et al. Deciphering the super relaxed state of human β-cardiac myosin and the mode of action of mavacamten from myosin molecules to muscle fibers. Proc. Natl. Acad. Sci. U. S. A. 201809540 (2018) doi:10.1073/pnas.1809540115.

24. Planelles-Herrero, V. J., Hartman, J. J., Robert-Paganin, J., Malik, F. I. & Houdusse, A. Mechanistic and structural basis for activation of cardiac myosin force production by omecamtiv mecarbil. Nat. Commun. 8, 190 (2017).

25. Houdusse, a, Szent-Gyorgyi, a G. & Cohen, C. Three conformational states of scallop myosin S1. Proc. Natl. Acad. Sci. 97, 11238–11243 (2000).

26. Wilson, C., Naber, N., Pate, E. & Cooke, R. The Myosin Inhibitor Blebbistatin Stabilizes the Super-Relaxed State in Skeletal Muscle. Biophys. J. 107, 1637–1646 (2014).

27. Alamo, L. et al. Lessons from a tarantula: new insights into muscle thick filament and myosin interacting-heads motif structure and function. Biophys. Rev. 9, 461–480 (2017).

28. Nag, S. et al. The myosin mesa and the basis of hypercontractility caused by hypertrophic cardiomyopathy mutations. Nat. Struct. Mol. Biol. 24, 525–533 (2017).

29. Coureux, P.-D., Sweeney, H. L. & Houdusse, A. Three myosin V structures delineate essential features of chemo-mechanical transduction. EMBO J. 23, 4527–4537 (2004).

30. Pylypenko, O. & Houdusse, A. M. Essential ‘ankle’ in the myosin lever arm. Proc. Natl. Acad. Sci. U. S. A. 108, 5–6 (2011).

31. Craig, R. & Padrón, R. Structural basis of the super-and hyper-relaxed states of myosin II. J. Gen. Physiol. 154, (2022).

32. Ikebe, M., Reardon, S., Schwonek, J. P., Sanders, C. R. 2nd & Ikebe, R. Structural requirement of the regulatory light chain of smooth muscle myosin as a substrate for myosin light chain kinase. J. Biol. Chem. 269, 28165–28172 (1994).

33. Toepfer, C. et al. Myosin regulatory light chain (RLC) phosphorylation change as a modulator of cardiac muscle contraction in disease. J. Biol. Chem. 288, 13446–13454 (2013).

34. Padrón, R. et al. The myosin interacting-heads motif present in live tarantula muscle explains tetanic and posttetanic phosphorylation mechanisms. Proc. Natl. Acad. Sci. U. S. A. 117, 11865–11874 (2020).

35. Huxley, H. E. & Brown, W. The low-angle x-ray diagram of vertebrate striated muscle and its behaviour during contraction and rigor. J. Mol. Biol. 30, 383–434 (1967).

36. Luther, P. K. et al. Understanding the organisation and role of myosin binding protein C in normal striated muscle by comparison with MyBP-C knockout cardiac muscle. J. Mol. Biol. 384, 60–72 (2008).

37. Al-Khayat, H. A., Morris, E. P., Kensler, R. W. & Squire, J. M. Myosin filament 3D structure in mammalian cardiac muscle. J. Struct. Biol. 163, 117–126 (2008).

38. Brito, R. et al. A molecular model of phosphorylation-based activation and potentiation of tarantula muscle thick filaments. J. Mol. Biol. 414, 44–61 (2011).

39. Hernandez, O. M., Jones, M., Guzman, G. & Szczesna-Cordary, D. Myosin essential light chain in health and disease. Am. J. Physiol. Heart Circ. Physiol. 292, H1643–54 (2007).

40. Toepfer, C. N. et al. Myosin Sequestration Regulates Sarcomere Function, Cardiomyocyte Energetics, and Metabolism, Informing the Pathogenesis of Hypertrophic Cardiomyopathy. Circulation 141, 828–842 (2020).

41. Sarkar, S. S. et al. The hypertrophic cardiomyopathy mutations R403Q and R663H increase the number of myosin heads available to interact with actin. Sci. Adv. 6, eaax0069 (2020).

42. Vander Roest, A. S., et al. Hypertrophic cardiomyopathy β-cardiac myosin mutation (P710R) leads to hypercontractility by disrupting super relaxed state. Proc. Natl. Acad. Sci. U. S. A. 118, (2021).

43. Huang, W. & Szczesna-Cordary, D. Molecular mechanisms of cardiomyopathy phenotypes associated with myosin light chain mutations. J. Muscle Res. Cell Motil. 36, 433–445 (2015).

44. Sitbon, Y. H. et al. Cardiomyopathic mutations in essential light chain reveal mechanisms regulating the super relaxed state of myosin. J. Gen. Physiol. 153, (2021).

45. Robert-Paganin, J. et al. The actomyosin interface contains an evolutionary conserved core and an ancillary interface involved in specificity. Nat. Commun. 12, 1892 (2021).

46. Haraguchi, T. et al. Discovery of ultrafast myosin, its amino acid sequence, and structural features. Proc. Natl. Acad. Sci. U. S. A. 119, (2022).

47. Ho, C. Y. et al. Genotype and Lifetime Burden of Disease in Hypertrophic Cardiomyopathy: Insights from the Sarcomeric Human Cardiomyopathy Registry (SHaRe). Circulation 138, 1387–1398 (2018).

48. McNamara, J. W. et al. Ablation of cardiac myosin binding protein-C disrupts the super-relaxed state of myosin in murine cardiomyocytes. J. Mol. Cell. Cardiol. 94, 65–71 (2016).

49. Burley, S. K. Integrative/Hybrid Methods Structural Biology: Role of Macromolecular Crystallography. Adv. Exp. Med. Biol. 1105, 11–18 (2018).

50. Patwardhan, A. & Lawson, C. L. Databases and Archiving for CryoEM. Methods Enzymol. 579, 393–412 (2016).

## References

1. Heissler, S. M., Arora, A. S., Billington, N., Sellers, J. R. & Chinthalapudi, K. Cryo-EM structure of the autoinhibited state of myosin-2. Sci. Adv. 7, eabk3273 (2021).

2. Planelles-Herrero, V. J., Hartman, J. J., Robert-Paganin, J., Malik, F. I. & Houdusse, A. Mechanistic and structural basis for activation of cardiac myosin force production by omecamtiv mecarbil. Nat. Commun. 8, 190 (2017).

3. Robert-Paganin, J., Auguin, D. & Houdusse, A. Hypertrophic cardiomyopathy disease results from disparate impairments of cardiac myosin function and auto-inhibition. Nat. Commun. 9, 4019 (2018).

4. Nag, S. et al. The myosin mesa and the basis of hypercontractility caused by hypertrophic cardiomyopathy mutations. Nat. Struct. Mol. Biol. 24, 525–533 (2017).

5. Spudich, J. A. Three perspectives on the molecular basis of hypercontractility caused by hypertrophic cardiomyopathy mutations. Pflugers Arch. 471, 701–717 (2019).

6. Brito, R. et al. A molecular model of phosphorylation-based activation and potentiation of tarantula muscle thick filaments. J. Mol. Biol. 414, 44–61 (2011).

7. Rasicci, D. V et al. Dilated cardiomyopathy mutation E525K in human beta-cardiac myosin stabilizes the interacting heads motif and super-relaxed state of myosin. bioRxiv 2022.02.18.480995 (2022) doi:10.1101/2022.02.18.480995.

8. Kandiah, E. et al. CM01: a facility for cryo-electron microscopy at the European Synchrotron. Acta Crystallogr. Sect. D 75, 528–535 (2019).

9. Zheng, S. Q. et al. MotionCor2: anisotropic correction of beam-induced motion for improved cryo-electron microscopy. Nat. Methods 14, 331–332 (2017).

10. Zhang, K. Gctf: Real-time CTF determination and correction. J. Struct. Biol. 193, 1–12 (2016).

11. Punjani, A., Rubinstein, J. L., Fleet, D. J. & Brubaker, M. A. cryoSPARC: algorithms for rapid unsupervised cryo-EM structure determination. Nat. Methods 14, 290–296 (2017).

12. Scheres, S. H. W. & Chen, S. Prevention of overfitting in cryo-EM structure determination. Nature methods vol. 9 853–854 (2012).

13. Blankenfeldt, W., Thomä, N. H., Wray, J. S., Gautel, M. & Schlichting, I. Crystal structures of human cardiac beta-myosin II S2-Delta provide insight into the functional role of the S2 subfragment. Proc. Natl. Acad. Sci. U. S. A. 103, 17713–17717 (2006).

14. Brooks, B. R. et al. CHARMM: The biomolecular simulation program. J. Comput. Chem. 30, 1545–1614 (2009).

15. Jo, S., Kim, T., Iyer, V. G. & Im, W. CHARMM-GUI: A web-based graphical user interface for CHARMM. J. Comput. Chem. 29, 1859–1865 (2008).

16. Liebschner, D. et al. Macromolecular structure determination using X-rays, neutrons and electrons: recent developments in Phenix. *Acta Crystallogr. Sect. D, Struct*. Biol. 75, 861–877 (2019).

17. Kovalevskiy, O., Nicholls, R. A., Long, F., Carlon, A. & Murshudov, G. N. Overview of refinement procedures within REFMAC5: utilizing data from different sources. *Acta Crystallogr. Sect. D, Struct*. Biol. 74, 215–227 (2018).

18. Huang, J. et al. CHARMM36m: an improved force field for folded and intrinsically disordered proteins. Nat. Methods 14, 71–73 (2017).

19. Abraham, M. J. et al. Gromacs: High performance molecular simulations through multi-level parallelism from laptops to supercomputers. SoftwareX 1–2, 19–25 (2015).

20. Chen, V. B. et al. MolProbity: all-atom structure validation for macromolecular crystallography. Acta Crystallogr. D. Biol. Crystallogr. 66, 12–21 (2010).

21. Barad, B. A. et al. EMRinger: side chain-directed model and map validation for 3D cryo-electron microscopy. Nat. Methods 12, 943–946 (2015).

22. Laskowski, R. A. PDBsum new things. Nucleic Acids Res. 37, D355–9 (2009).

23. Sarkar, S. S. et al. The hypertrophic cardiomyopathy mutations R403Q and R663H increase the number of myosin heads available to interact with actin. Sci. Adv. 6, eaax0069 (2020).

24. Charron, P. et al. Prenatal molecular diagnosis in hypertrophic cardiomyopathy: report of the first case. Prenat. Diagn. 24, 701–703 (2004).

25. Keller, D. I. et al. Human homozygous R403W mutant cardiac myosin presents disproportionate enhancement of mechanical and enzymatic properties. J. Mol. Cell. Cardiol. 36, 355–362 (2004).

26. Alamo, L., Pinto, A., Sulbarán, G., Mavárez, J. & Padrón, R. Lessons from a tarantula: new insights into myosin interacting-heads motif evolution and its implications on disease. Biophys. Rev. (2017) doi:10.1007/s12551-017-0292-4.

27. Bos, J. M. et al. Characterization of a Phenotype-Based Genetic Test Prediction Score for Unrelated Patients With Hypertrophic Cardiomyopathy. Mayo Clin. Proc. 89, 727–737 (2014).

28. Bloemink, M. et al. The Hypertrophic Cardiomyopathy Myosin Mutation R453C Alters ATP Binding and Hydrolysis of Human Cardiac β-Myosin. J. Biol. Chem. 289, 5158–5167 (2014).

29. Sommese, R. F. et al. Molecular consequences of the R453C hypertrophic cardiomyopathy mutation on human -cardiac myosin motor function. Proc. Natl. Acad. Sci. 110, 12607–12612 (2013).

30. Spudich, J. A. et al. Effects of hypertrophic and dilated cardiomyopathy mutations on power output by human β-cardiac myosin. J. Exp. Biol. 219, 161–7 (2016).

31. Yu, C.-M. et al. Left ventricular reverse remodeling but not clinical improvement predicts long-term survival after cardiac resynchronization therapy. Circulation 112, 1580–1586 (2005).

32. Pablo, K. J. et al. Prevalence of Sarcomere Protein Gene Mutations in Preadolescent Children With Hypertrophic Cardiomyopathy. Circ. Cardiovasc. Genet. 2, 436–441 (2009).

33. Adhikari, A. S. et al. Early-Onset Hypertrophic Cardiomyopathy Mutations Significantly Increase the Velocity, Force, and Actin-Activated ATPase Activity of Human β-Cardiac Myosin. Cell Rep. 17, 2857–2864 (2016).

34. Adhikari, A. S. et al. β-Cardiac myosin hypertrophic cardiomyopathy mutations release sequestered heads and increase enzymatic activity. Nat. Commun. 10, 2685 (2019).

35. Kawana, M., Sarkar, S. S., Sutton, S., Ruppel, K. M. & Spudich, J. A. Biophysical properties of human beta-cardiac myosin with converter mutations that cause hypertrophic cardiomyopathy. Sci Adv 3, e1601959 (2017).

36. Kuang, S. Q. et al. Identification of a novel missense mutation in the cardiac beta-myosin heavy chain gene in a Chinese patient with sporadic hypertrophic cardiomyopathy. J. Mol. Cell. Cardiol. 28, 1879–1883 (1996).

37. Millat, G. et al. Prevalence and spectrum of mutations in a cohort of 192 unrelated patients with hypertrophic cardiomyopathy. Eur. J. Med. Genet. 53, 261–267 (2010).

38. Kassem, H. S. et al. Early Results of Sarcomeric Gene Screening from the Egyptian National BA-HCM Program. J. Cardiovasc. Transl. Res. 6, 65–80 (2013).

39. Nanni, L. et al. Hypertrophic cardiomyopathy: two homozygous cases with ‘typical’ hypertrophic cardiomyopathy and three new mutations in cases with progression to dilated cardiomyopathy. Biochem. Biophys. Res. Commun. 309, 391–8 (2003).

40. Song, L. et al. Mutations profile in Chinese patients with hypertrophic cardiomyopathy. Clin. Chim. Acta 351, 209–216 (2005).

41. Olson, T. M., Karst, M. L., Whitby, F. G. & Driscoll, D. J. Myosin light chain mutation causes autosomal recessive cardiomyopathy with mid-cavitary hypertrophy and restrictive physiology. Circulation 105, 2337–2340 (2002).

42. Sitbon, Y. H. et al. Cardiomyopathic mutations in essential light chain reveal mechanisms regulating the super relaxed state of myosin. J. Gen. Physiol. 153, (2021).

43. Yadav, S., Sitbon, Y. H., Kazmierczak, K. & Szczesna-Cordary, D. Hereditary heart disease: pathophysiology, clinical presentation, and animal models of HCM, RCM, and DCM associated with mutations in cardiac myosin light chains. Pflugers Arch. 471, 683–699 (2019).

44. Mohiddin, S. A. et al. Utility of genetic screening in hypertrophic cardiomyopathy: prevalence and significance of novel and double (homozygous and heterozygous) beta-myosin mutations. Genet. Test. 7, 21–27 (2003).

45. Huang, W. & Szczesna-Cordary, D. Molecular mechanisms of cardiomyopathy phenotypes associated with myosin light chain mutations. J. Muscle Res. Cell Motil. 36, 433–445 (2015).

46. Robert-Paganin, J., Auguin, D. & Houdusse, A. Hypertrophic cardiomyopathy disease results from disparate impairments of cardiac myosin function and auto-inhibition. Nat. Commun. 9, 4019 (2018).

47. N., T. C., et al. Myosin Sequestration Regulates Sarcomere Function, Cardiomyocyte Energetics, and Metabolism, Informing the Pathogenesis of Hypertrophic Cardiomyopathy. Circulation 141, 828–842 (2020).

48. Vander Roest, A. S., et al. Hypertrophic cardiomyopathy β-cardiac myosin mutation (P710R) leads to hypercontractility by disrupting super relaxed state. Proc. Natl. Acad. Sci. U. S. A. 118, (2021).

49. Alamo, L. et al. Lessons from a tarantula: new insights into muscle thick filament and myosin interacting-heads motif structure and function. Biophys. Rev. 9, 461–480 (2017).

50. Risi, C. et al. High-Resolution Cryo-EM Structure of the Cardiac Actomyosin Complex. Structure 29, 50–60.e4 (2021).

51. Nag, S. et al. Contractility parameters of human -cardiac myosin with the hypertrophic cardiomyopathy mutation R403Q show loss of motor function. Sci. Adv. 1, e1500511– e1500511 (2015).

52. Vera, C. D. et al. Myosin motor domains carrying mutations implicated in early or late onset hypertrophic cardiomyopathy have similar properties. J. Biol. Chem. 294, 17451–17462 (2019).

53. Adhikari, A. S. et al. Early-Onset Hypertrophic Cardiomyopathy Mutations Significantly Increase the Velocity, Force, and Actin-Activated ATPase Activity of Human beta-Cardiac Myosin. Cell Rep 17, 2857–2864 (2016).

54. Bell, K. M., Kronert, W. A., Huang, A., Bernstein, S. I. & Swank, D. M. The R249Q hypertrophic cardiomyopathy myosin mutation decreases contractility in Drosophila by impeding force production. J. Physiol. 597, 2403–2420 (2019).

55. Kawana, M., Sarkar, S. S., Sutton, S., Ruppel, K. M. & Spudich, J. Biophysical properties of human β-cardiac myosin with converter mutations that cause hypertrophic cardiomyopathy. Sci Adv 3, 1–11 (2016).

56. Robert-Paganin, J., Pylypenko, O., Kikuti, C., Sweeney, H. L. & Houdusse, A. Force Generation by Myosin Motors: A Structural Perspective. Chem. Rev. 120, 5–35 (2020).

57. Fourey, D. et al. Prevalence and Clinical Implication of Double Mutations in Hypertrophic Cardiomyopathy: Revisiting the Gene-Dose Effect. Circ. Cardiovasc. Genet. 10, (2017).

58. Morck, M. M. et al. Hypertrophic cardiomyopathy mutations in the pliant and light chain-binding regions of the lever arm of human β-cardiac myosin have divergent effects on myosin function. Elife 11, e76805 (2022).

59. Singh, R. R., McNamara, J. W. & Sadayappan, S. Mutations in myosin S2 alter cardiac myosin-binding protein-C interaction in hypertrophic cardiomyopathy in a phosphorylation-dependent manner. J. Biol. Chem. 297, 100836 (2021).

